# Functional remodeling of lysosomes by type I interferon modifies host defense

**DOI:** 10.1101/2020.02.25.965061

**Authors:** Hailong Zhang, Abdelrahim Zoued, Xu Liu, Brandon Sit, Matthew K. Waldor

## Abstract

Organelle remodeling is critical for cellular homeostasis, but host factors that control organelle function during microbial infection remain largely uncharacterized. Here, a genome-scale CRISPR/Cas9 screen in intestinal epithelial cells with the prototypical intracellular bacterial pathogen *Salmonella* led us to discover that type I interferon (IFN-I) remodels lysosomes. Even in the absence of infection, IFN-I signaling modified the localization, acidification, protease activity and proteomic profile of lysosomes. Proteomic and genetic analyses revealed that multiple IFN-I-stimulated genes including *Ifitm3, Slc15a3*, and *Cnp* contribute to lysosome acidification. IFN-I-dependent lysosome acidification stimulated intracellular *Salmonella* virulence gene expression, leading to rupture of the *Salmonella*-containing vacuole and host cell death. Moreover, IFN-I signaling promoted *in vivo Salmonella* pathogenesis in the intestinal epithelium, where *Salmonella* initiates infection. Our findings explain how an intracellular bacterial pathogen co-opts epithelial IFN-I signaling. We propose that IFN-I control of lysosome function broadly impacts host defense against diverse viral and microbial pathogens.

## INTRODUCTION

Microbial pathogens have evolved varied virulence strategies to modulate host cell function (Geoghegan and Holmes, 2018; Ribet and Cossart, 2015). A common mechanism, employed by all viral and some bacterial pathogens, is to enter host cells, where they co-opt cellular functions while simultaneously evading extracellular threats such as innate and adaptive immune mechanisms (Hybiske and Stephens, 2008; Lee et al., 2019). Inside cells, intracellular pathogens interact with and exploit host cell organelles to support their proliferation (Omotade and Roy, 2019). As a result of their intimate relationships with and manipulation of varied host cell functions, intracellular pathogens have proven to be outstanding tools to probe basic eukaryotic cell biology (Welch, 2015).

Compared to knowledge of how microbial pathogens interact with phagocytic cells, less is known about the landscape of pathogen-epithelial cell interactions at barrier sites, where most infections originate (Jo, 2019). The human foodborne pathogen *Salmonella enterica* serovar Typhimurium (Stm) is a model intracellular bacterium that initially invades and subsequently kills intestinal epithelial cells (IECs) before spreading systemically via circulating phagocytes (Hurley et al., 2014). Stm’s entry into and initial trafficking inside IEC is well-characterized, and a hallmark of Stm infection is the formation of the *Salmonella*-containing vacuole (SCV), a dynamic, lysosome-like compartment that is permissive for Stm replication (Steele-Mortimer, 2008; Tuli and Sharma, 2019). However, the IEC pathways that control Stm-induced cytotoxicity remain incompletely defined.

One striking feature of the host response to Stm is the induction of type I interferons (IFN-Is), which include IFNα and IFNβ (Hess et al., 1989). IFN-Is are cytokines that, once secreted, bind the IFN-I receptor (IFNAR1/2) to activate JAK-STAT signaling, which triggers expression of intracellular anti-microbial transcriptional programs consisting of over 400 IFN-stimulated genes (ISGs) (Schoggins and Rice, 2011). Due to the large size of the “interferome” and the complex interactions of ISGs with thousands of additional cellular proteins (Hubel et al., 2019), knowledge of the full spectrum of IFN-I-mediated changes in cellular function is incomplete. Although IFN-Is are known to have critical roles in antiviral responses, their functions in bacterial infection are less clear, and IFN-I signaling has been reported to be either protective or detrimental to the host depending on the specific bacterial pathogen (Kovarik et al., 2016).

Here, we carried out a genome-scale CRISPR/Cas9 screen to identify the host factors that contribute to Stm’s cytotoxicity to IECs. This screen revealed IFN-I signaling as a key susceptibility factor for cytotoxicity in IECs and led to our discovery of a novel role for IFN-I signaling in lysosomal localization and function, including the modification of this organelle’s pH, protease activity, and protein content. Organellar proteomics revealed that 11 ISGs were enriched in lysosomes following IFN-I stimulation, several of which were found to directly impact lysosomal pH. IFN-I signaling-dependent lysosomal acidification stimulated Stm virulence gene expression, and *in vivo* studies confirmed a role for epithelial IFN-I signaling in promoting systemic Stm infection. IFN-I signaling mediated control of lysosome function likely contributes to host responses to diverse intracellular pathogens and viruses.

## RESULTS

### A CRISPR/Cas9 screen identifies IEC factors required for Stm cytotoxicity

As large-scale genetic screens to identify epithelial cell factors that mediate interactions with intracellular pathogens are lacking, we performed a multi-round, genome-wide CRISPR/Cas9 loss-of-function screen in the human colonic epithelial cell line HT29-Cas9 (Blondel et al., 2016), to identify IEC genes that confer resistance to Stm cytotoxicity (Figure 1A). HT29 cells are efficiently invaded and subsequently killed by Stm, providing a strong selective force to enrich for guide RNAs targeting host factors that modulate cytotoxicity. The screen identified known pathways that sensitize cells to Stm infection, including those involved in regulation of actin dynamics (Arp2/3 and Rac), which are important in pathogen invasion (Table S1 and Figures 1B, C), (Unsworth et al., 2004; Yeung et al., 2019). Genes linked to pathways not previously directly linked to Stm virulence, including the Fc-gamma receptor-dependent phagocytic and GPI anchor modification pathways were also enriched (Figures 1B, C). Strikingly, the top ‘hits’ of the screen were remarkably coherent - the seven most enriched guide RNAs from both libraries screened corresponded to genes in the IFN-I signaling pathway, including the receptor (*Ifnar1*/*Ifnar2*), adaptor (*Jak1*/*Tyk2*), and transcription factor (*Stat1*/*Stat2*/*Irf9*) components of the system (Figures 1D, E), suggesting that IFN-I signaling is a major driver of Stm-mediated cytotoxicity in IECs.

**Figure 1.**
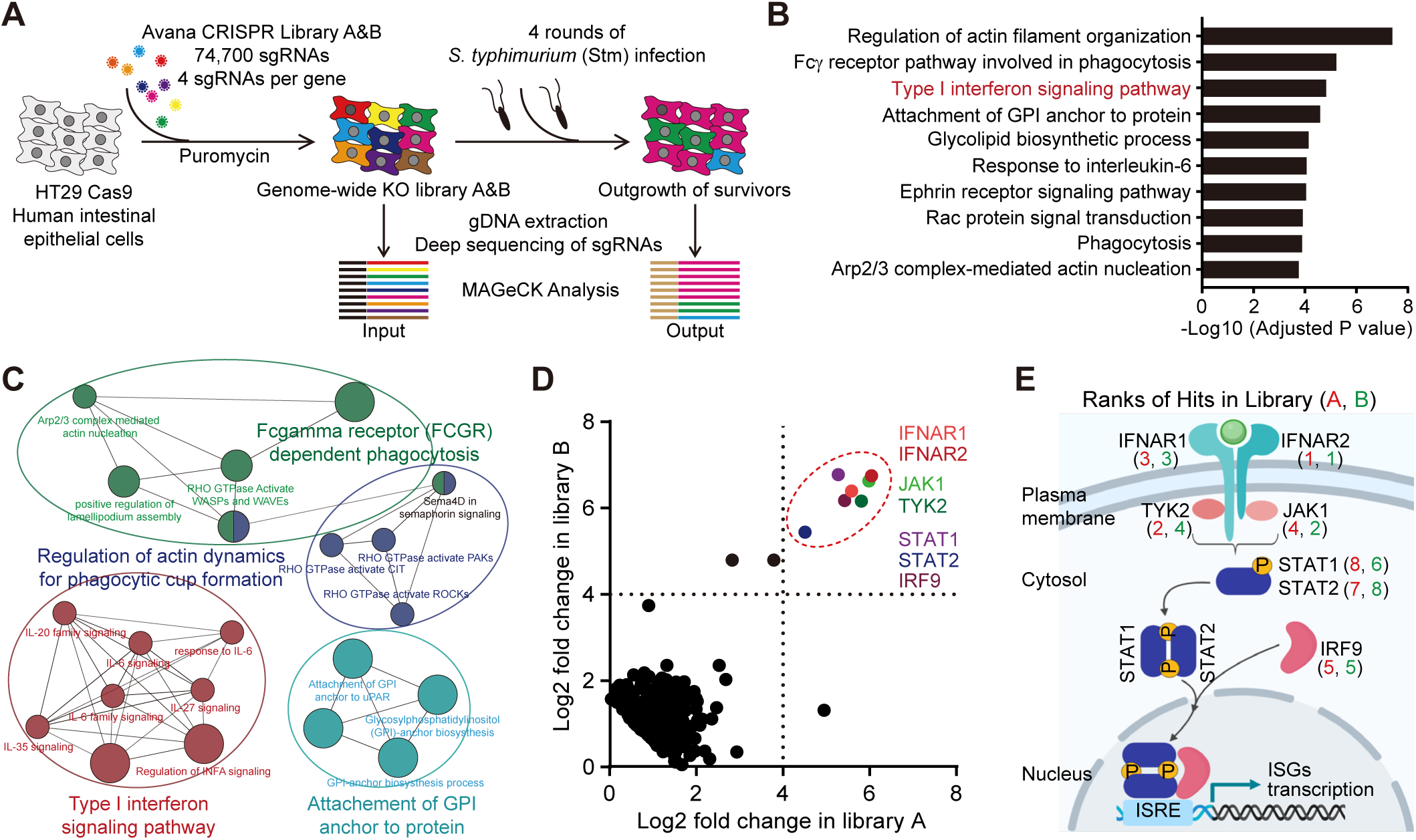
A CRISPR/Cas9 screen identifies IEC factors required for Stm cytotoxicity. (A) Workflow for CRISPR/Cas9 Stm cytotoxicity screen in HT29-Cas9 cells. (B) Adjusted p-values for selected enriched Gene Ontology (GO) terms from GO-analyzed hits in the Stm cytotoxicity screen (upper threshold set at p < 1E-03). (C) Cytoscape visualization of enriched pathways. (D) Scatterplots showing normalized read enrichment of specific sgRNAs in two libraries (A and B) after 4 rounds of Stm infection. Genes involved in IFN-I signaling are delineated by the dashed red circle. (E) Overview of IFN-I signaling pathway. Numbers correspond to hit ranks in each library.

### IFN-I promotes Stm cytotoxicity in IECs

Stm induces IFN-I production during infection (Hess et al., 1989), but the function of IFN-I signaling in IECs is unknown. A clonal knockout of *Ifnar2*, the top enriched hit in both libraries, was constructed in the HT29-Cas9 background (Figure S1A), to validate the screen findings. At both early and late infection time points, *Ifnar2* KO cells were more resistant to Stm-induced cell death than the wild-type (WT) parental line (Figure 2A and S1B, C). Priming cells with IFNβ (a major IFN-I), conditions that mimic the multiple rounds of the original screen, further sensitized WT but not *Ifnar2* KO cells to death (Figure 2A). To complement these findings, we treated WT cells with chemical inhibitors of JAK-STAT signaling, the downstream target of activated IFNAR1/2. Similar to *Ifnar2* KO cells, treatment with the JAK inhibitors ruxolitinib and pyridone-6 also diminished Stm*-*induced death in WT cells, indicating active IFN-I signaling is required for this phenotype (Figure 2B).

**Figure 2.**
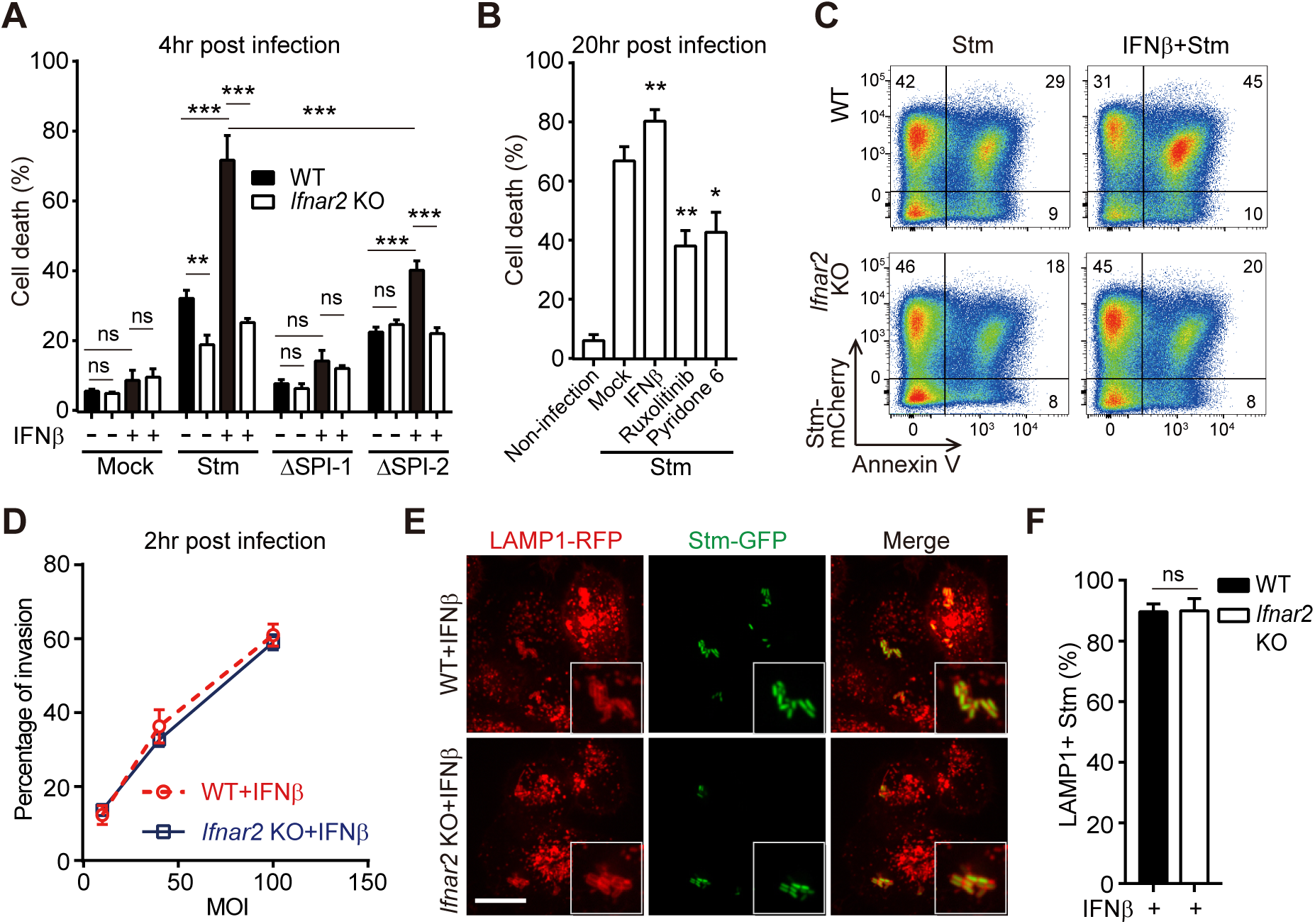
IFN-I promotes Stm cytotoxicity in IECs. (A) Survival of IFNβ-primed or unprimed WT or *Ifnar2* KO HT29 cells 4 hours post WT or mutant Stm infection. Mean ± s.d., n = 3. (B) Survival of mock or drug-treated WT HT29 cells 20 hours post WT Stm infection. Mean ± s.d., n = 3. (C) Flow cytometry of IFNβ-primed or unprimed WT and *Ifnar2* KO HT29 cells 20 hours post mCherry-Stm infection and stained with Annexin V-FITC. FITC, fluorescein isothiocyanate. (D) Flow cytometric quantification of invasion of HT29 cells by mCherry-Stm. Mean ± s.d., n = 3. (E) Representative images of LAMP1-RFP-expressing HeLa cells 4 hours post GFP-Stm infection. Boxed insets depict higher magnification showing bacterial colocalization with LAMP1-RFP. Scale bar, 10μm. (F) Quantification of LAMP1-RFP-positive Stm from 10 fields. Mean ± s.d., n = 3. Statistical analysis was performed by two-tailed Student’s *t* test (*P < 0.05, **P < 0.01 and ***P < 0.001). See also Figure S1, 2 and Table S1.

Our overall observations that IFN-I promotes Stm*-*induced IEC death is consistent with prior data that this cytokine enhances necroptosis in Stm*-*infected macrophages (Robinson et al., 2012). However, it is not clear whether macrophage and epithelial cell responses to Stm infection are similar; furthermore, it is known that Stm*-*induced cell death in macrophages is invasion-independent (van der Velden et al., 2000). Thus, we next tested whether IFN-I-promoted epithelial cell death depended on SPI-1 or SPI-2, critical *Salmonella* pathogenicity islands that each encode type 3 secretion systems (T3SS) required for cellular invasion and intracellular survival, respectively (Galan et al., 2014). SPI-1-deficient (Δ*prgH*) Stm did not induce epithelial cell death in any condition, confirming that in IECs cytotoxicity requires cell invasion (Figure 2A). In contrast, SPI-2-deficient (Δ*ssaV*) Stm led to reduced but still detectable levels of cytotoxicity that remained sensitive to IFNβ priming, suggestive of both SPI-2-dependent and independent mechanisms of intracellular Stm-induced cytotoxicity (Figure 2A).

In support of the population-level LDH assays, flow cytometry of HT29 or HeLa cells infected with fluorescent Stm and stained with the cell death probe Annexin-V indicated that IFN-I only influenced cell death in the population of cells that contained intracellular Stm (Figures 2C and S1D-F). In addition, we found that. IFN-I signaling did not impact Stm invasion (Figures 2D and S2A), nor did it influence bacterial association with the early endosomal marker Rab5, late endosomal marker Rab7 or lysosomal marker LAMP1 (Desjardins et al., 1994) (Figures 2E, F and S2B-E). Together, these data suggest that IFN-I-mediated sensitization of epithelial cells to Stm occurs downstream of cell invasion and initial SCV formation.

### IFN-I regulates lysosome localization and function

During our analyses of SCV formation, we unexpectedly observed that IFN-I signaling alters the subcellular localization of lysosomes in epithelial cells, even in the absence of infection. In HeLa cells, lysosomes (identifiable as LAMP1+/Lysotracker+ co-staining organelles) were scattered throughout the cytoplasm under basal conditions. Following IFNβ stimulation, lysosomes re-localized to the perinuclear region (Figures 3A, B, Video S1, 2); lysosome re-localization was not observed in *Ifnar2* KO HeLa cells, confirming that this response was dependent on IFN-I signaling.

**Figure 3.**
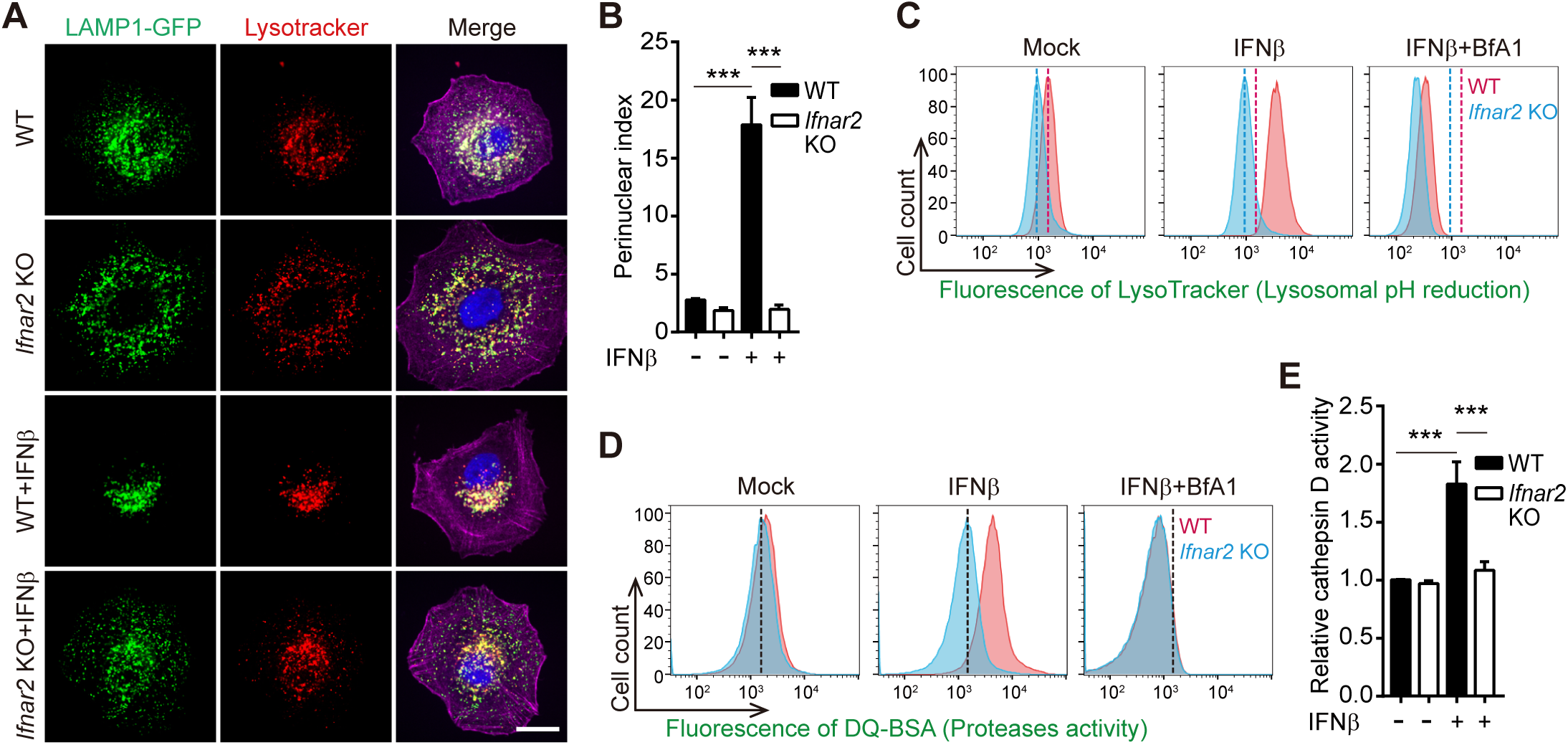
IFN-I signaling regulates lysosomal positioning, acidity, and protease activity. (A) Representative images of lysosome (LAMP1-GFP+/LysoTracker+ compartment) distribution in WT and *Ifnar2* KO HeLa cells with or without 16 hours of IFNβ stimulation. Nuclei (blue) were stained with DAPI and actin (purple) was stained with phalloidin. Scale bar, 5μm. (B) Quantification of perinuclear lysosome indices from 10 cells. Mean ± s.d., n = 3. (C-D) Flow cytometry of LysoTracker Red (C) and DQ-Green BSA fluorescence (D) in HeLa cells ±16 hours of treatment with IFNβ or the lysosomal acidification inhibitor Bfa1. Vertical dashed lines indicate the mean fluorescence value of the mock control in WT (red) or *Ifnar2* KO (blue) cells. (E) Relative cathepsin D activity in WT and *Ifnar2* KO HeLa cells ± 16 hours of IFNβ treatment. Mean ± s.d., n = 5. Statistical analysis was performed by two-tailed Student’s *t* test (***P < 0.001). See also Figure S3.

Notably, IFNβ priming led to significantly higher intensities of two fluorescent reporters (Lysotracker and Lysosensor) of lysosomal pH in WT but not *Ifnar2* KO cells, indicating that IFN-I signaling lowers lysosome pH (Figures 3C and S3A-C). Moreover, staining with fluorescent reporters of general lysosomal protease activity (DQ-BSA (Reis et al., 1998)), or cathepsin D (a major lysosomal protease) activity revealed that their activities were elevated by IFNβ stimulation in an *Ifnar2*-dependent fashion (Figures 3D, E and S3D). These findings are consistent with the idea that the activity of most resident lysosomal proteins, including cathepsins and other degradative enzymes, is positively regulated by acidic pH (Butor et al., 1995). Staining with fluorescent reporters of endocytic activity (Dextran-568) suggested that IFN-I signaling does not impact general endocytic trafficking; thus, IFN-I signaling appears to specifically influence lysosome function and positioning (Figure S3E). IFNβ-induced lysosomal acidification and protease activation was abolished by the addition of the v-ATPase inhibitor bafilomycin A1 (Bfa1) (Yoshimori et al., 1991) (Figure 3C, D), demonstrating that IFN-I signaling primarily relies on the conventional lysosomal acidification machinery.

Together, these observations reveal that IFN-I signaling promotes epithelial cell lysosomal re-localization, acidification and degradative activity, without broadly affecting intracellular trafficking. Besides epithelial cells such as HeLa and HT29, IFNβ treatment also reduced lysosomal pH in monocyte/macrophage-like THP-1 cells (Figure S3F), suggesting that IFN-I signaling controls lysosomal acidification outside of epithelial cell lineages.

### The ISG IFITM3 regulates lysosomal function and Stm cytotoxicity

To begin to dissect the mechanism by which IFN-I signaling regulates lysosome acidification and function, we first took a candidate-based approach and investigated IFITM3. This transmembrane ISG has antiviral activity and is thought to reside in the endosomal trafficking system and to interact with the lysosomal v-ATPase complex (Spence et al., 2019; Wee et al., 2012), suggesting a potential role for this protein in lysosome function. Immunofluorescence analysis revealed that IFITM3 co-localized with LAMP1, but not Rab5, confirming that IFITM3 is a lysosomal protein (Figure 4A). Remarkably, lysosomal pH in *Ifitm3* KO cells was elevated in both basal and IFNβ-primed conditions relative to WT cells, partially phenocopying the *Ifnar2* KO, and suggesting that IFITM3 contributes to IFN-I-meditated lysosomal remodeling (Figures 4B-D). *Ifitm3*’s contribution to basal pH levels are consistent with the tonic activities of IFNs observed in diverse mammalian cell types (Schoggins et al., 2014). *Ifitm3* KO cells were more resistant to Stm-induced cell death than the WT parental line both before and after IFNβ priming (Figure 4E), suggesting that this ISG contributes to IFN-I signaling augmentation of Stm cytotoxicity.

**Figure 4.**
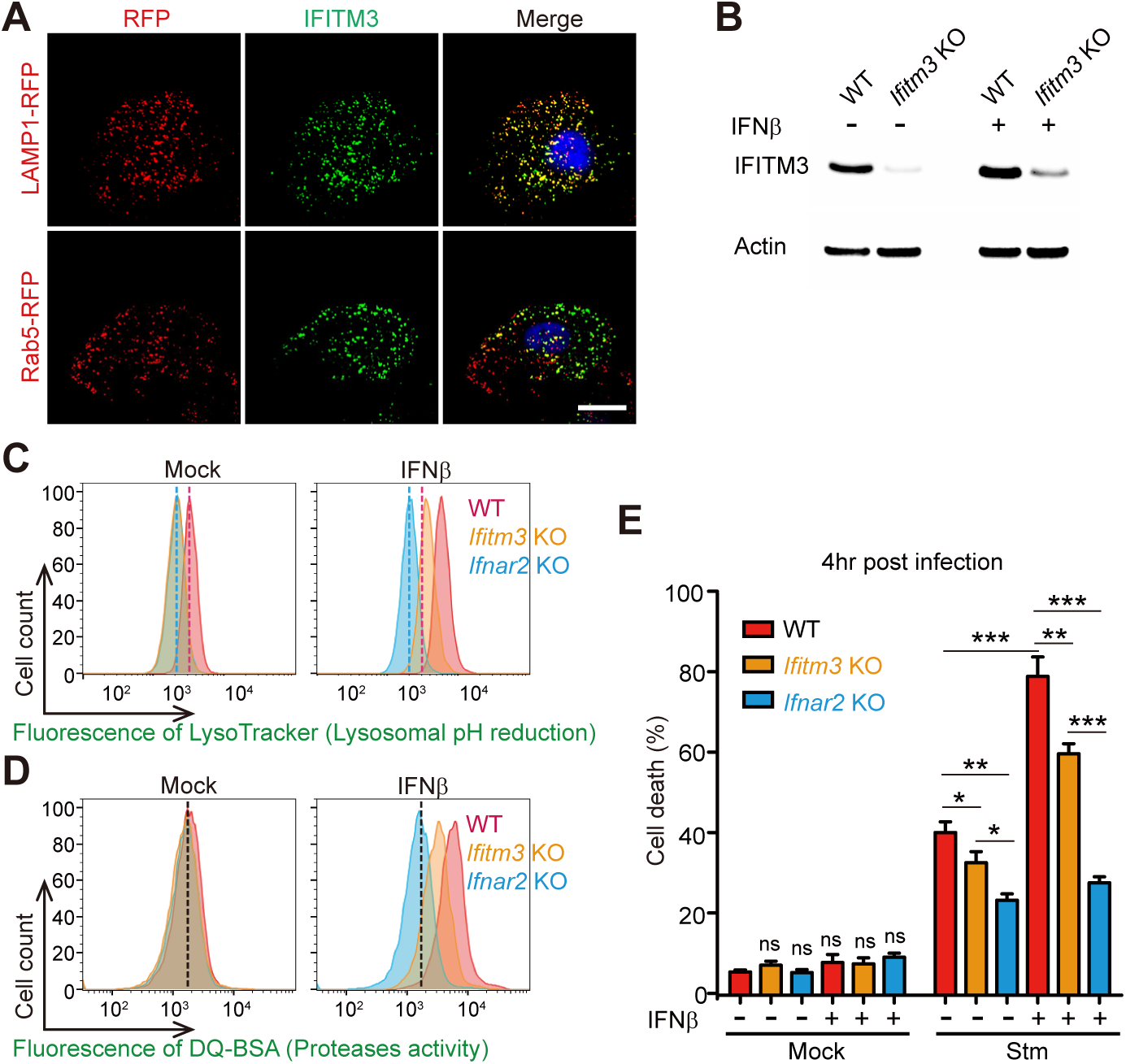
The ISG IFITM3 regulates lysosomal function and Stm cytotoxicity. (A) Representative images of LAMP1-RFP or Rab5-RFP–expressing HeLa cells stained with IFITM3 antibody (GFP). Nuclei (blue) were stained with DAPI. Scale bar, 5μm. (B) Immunoblotting for IFITM3 in WT and *Ifitm3* KO HeLa cells ± 16 hours of IFNβ treatment. (C-D) Flow cytometry of LysoTracker Red (D) and DQ-Green BSA fluorescence (E) in WT, *Ifitm3* KO and *Ifnar2* KO HeLa cells ± 16 hours of IFNβ treatment. (E) Survival of IFNβ-primed or unprimed WT, *Ifitm3* or *Ifnar2* KO HeLa cells 4 hours post Stm infection. Mean ± s.d., n = 3. Statistical analysis was performed by two-tailed Student’s *t* test (*P < 0.05, **P < 0.01 and ***P < 0.001).

### Discovery of ISGs with novel roles in lysosomal pH regulation

Given that both lysosomal pH and degradative activity in *Ifitm3* KO cells were still somewhat sensitive to IFNβ priming (Figures 4C, D), we hypothesized that additional ISGs regulate lysosome function. To identify these factors, we employed organellar proteomics, a powerful and unbiased affinity-based technique that has not yet been applied to host-pathogen interactions. We used a recently described lysosomal pulldown system, LysoIP (Abu-Remaileh et al., 2017), to profile the proteomes of intact lysosomes from WT or *Ifnar2* KO cells in basal or IFNβ-stimulated states. The purity and integrity of the lysosome samples was confirmed by verifying the presence of luminal cathepsin D and the absence of cytosolic and Golgi apparatus markers (Figures 5A and S4). Quantitative profiling revealed that the abundances of ∼15 proteins, most of them ISGs, were increased in purified lysosomes upon IFNβ stimulation (Figure 5B). Spectral counts for IFITM3 were enriched in lysosomes from IFNβ-treated cells, supporting the imaging above (Figure 4A) and providing validation of the dataset. Immunoblots of purified lysosomal and cytoplasmic fractions from naïve and IFNβ-treated cells with antibodies to IFITM3 further corroborated this observation, and immunoblots for the known cytosolic ISG IFIT3 served as a negative control in this assay (Figure 5A).

**Figure 5.**
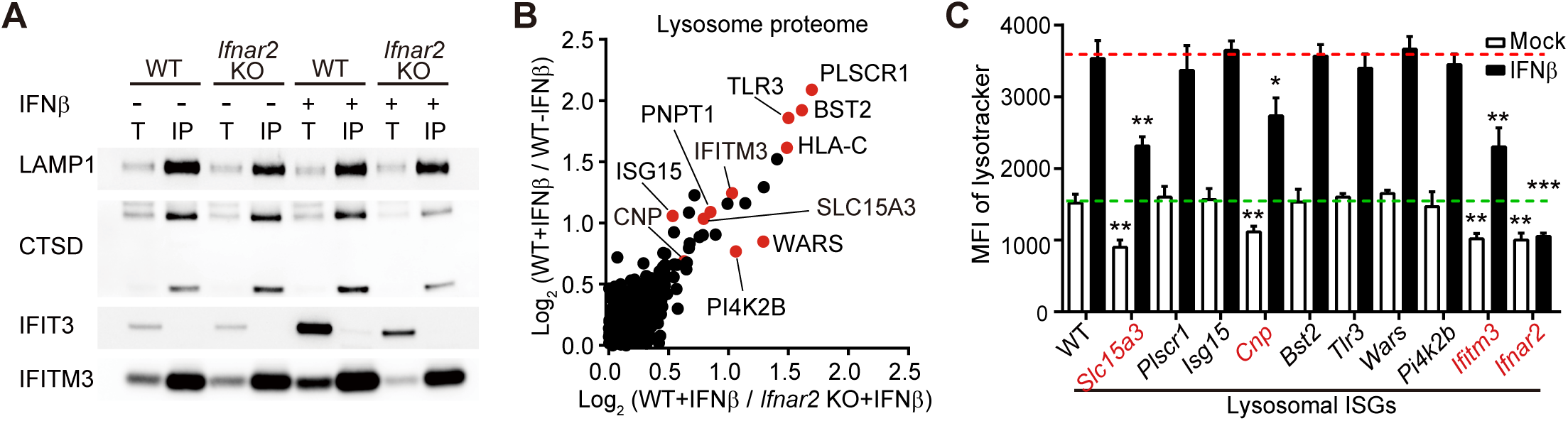
Discovery of ISGs with novel roles in lysosomal pH regulation. (A) Immunoblotting for known (LAMP1, CTSD) and suspected (IFITM3) lysosomal proteins in whole-cell lysates (T) and purified lysosomes (IP). (B) Relative fold change scatterplot of protein abundance in lysosomes purified from WT or *Ifnar2* KO HeLa cells ± 16 hours of IFNβ treatment. Colored dots indicate proteins that are known ISGs. (C) Quantification of mean fluorescence intensity from flow cytometry of Lysotracker staining in WT or ISG KO cells ± 16 hours of IFNβ treatment. Statistical analysis was performed by two-tailed Student’s *t* test (*P < 0.05, **P < 0.01 and ***P < 0.001). See also Figure S4 and Table S2.

We constructed KO cell lines for most of the lysosomally enriched ISGs and assessed their contributions to the pH of this degradative organelle (Figure 5C). Although most ISGs did not appear to influence lysosomal pH, we found two additional ISGs (*Slc15a3* and *Cnp*) that like *Ifitm3* contributed to both basal and IFN-I-mediated lysosomal acidification (Figure 5C). In contrast to *Ifitm3, Slc15a3*, a lysosome-resident, proton-coupled histidine and di-tripeptide transporter (Song et al., 2018), likely does not interact with the v-ATPase, suggesting that both v-ATPase-dependent and independent mechanisms may underlie IFN-I-induced lysosomal acidification. Similarly, *Cnp*, the other identified regulator of lysosome pH, is a 2’,3’-cyclic nucleotide 3’ phosphodiesterase whose activity has not been linked to the v-ATPase.

### IFN-I stimulates intracellular Stm virulence gene expression and facilitates SCV damage

We hypothesized that IFN-I’s role in lysosomal acidification might explain why IFN-I signaling enhances Stm cytotoxicity because acidic pH is known to stimulate expression of SPI-2-encoded and other virulence genes (Chakraborty et al., 2015; Prost et al., 2007). Consistent with this idea, IFNβ priming increased intracellular Stm SPI-2 encoded gene expression (Figures 6A and S5A). These genes were only induced after SCV formation (i.e. later than one-hour post-infection), and the effect of IFNβ priming was eliminated in *Ifnar2* KO cells. Furthermore, treatment with Bfa1 abolished IFNβ induction of SPI-2 expression (Figures 6A and S5A), suggesting that diminished SCV acidification is the primary mechanism of IFN-I-enhanced SPI-2 induction. Analyses of SPI-2 gene expression using flow cytometry and a fluorescent *P*_*sifB*_::*gfp* Stm reporter strain (Garmendia et al., 2003), confirmed this phenotype at single bacterial cell resolution (Figure 6B). Similar expression trends were also observed in known acid-induced, virulence-associated genes that are not encoded within SPI-2, such as *pagD* (Gunn et al., 1995) (Figures 6C and S5B, C). This is consistent with the observation above (Figure 2A) that SPI-2-deficient Stm retain some cytotoxicity. Importantly, the expression of SPI-1 genes, which encode invasion-specific functions, was not altered in infections with IFNβ priming or in *Ifnar2* KO cells (Figures 6D and S5D). Together, these data indicate that IFN-I-mediated acidification of lysosomes promotes intracellular Stm virulence gene expression.

**Figure 6.**
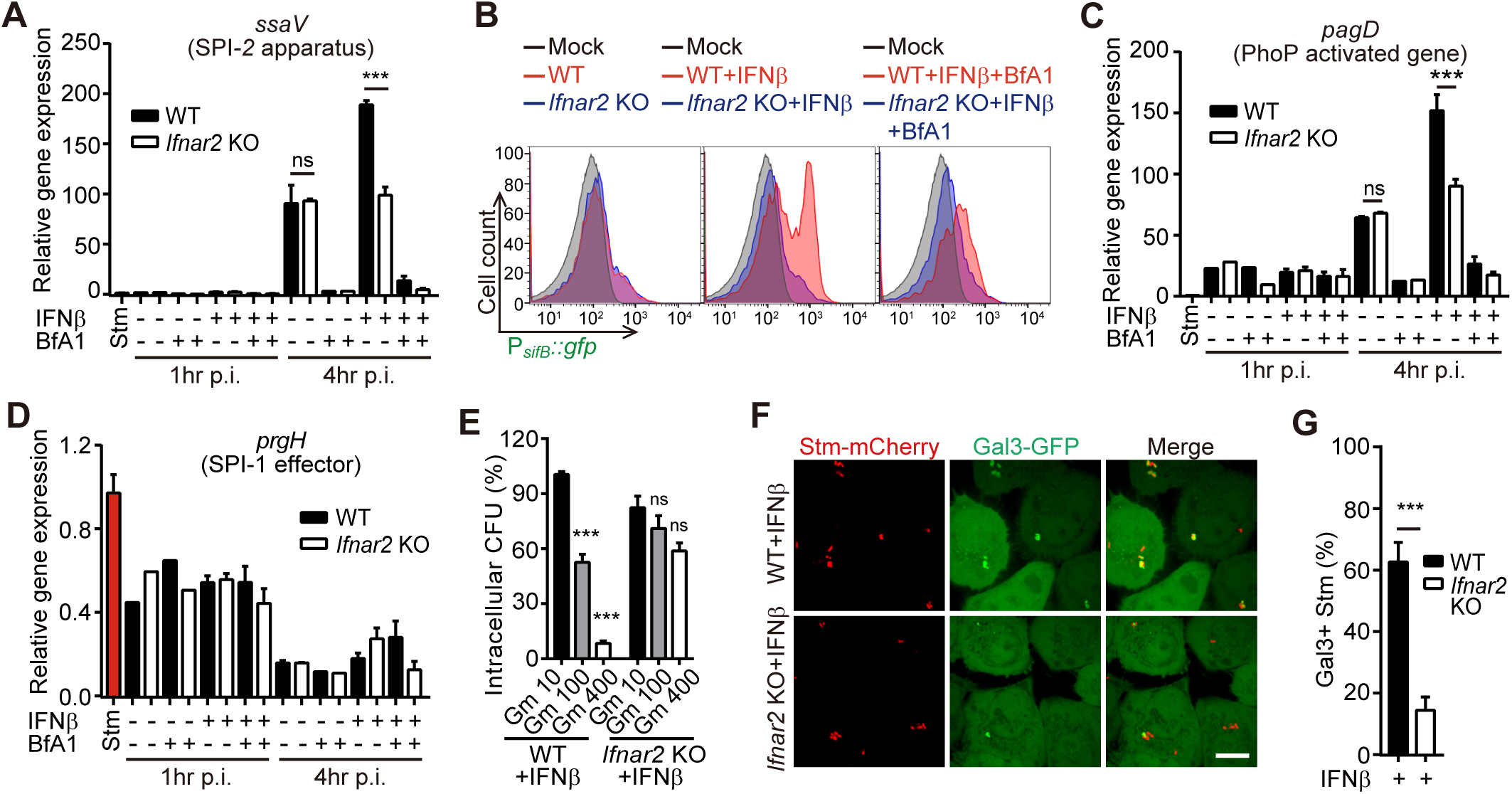
IFN-I signaling promotes Stm virulence gene expression and SCV rupture. Relative induction of SPI-2 (*ssaV*) (A), PhoP-induced virulence gene (*pagD*) (C) and SPI-1 (*prgH*) (D) in intracellular Stm from WT and *Ifnar2* KO HeLa cells ± 16 hours of drug treatment. Data are normalized to transcript levels in LB-cultured Stm (red). Mean ± s.d., n = 3. (B) Flow cytometry of intracellular P_*sifB*_::*gfp* Stm isolated from WT and *Ifnar2* KO HeLa cells ± 16 hours of drug treatment. LB-cultured Stm were used as the mock control. (E) Intracellular CFU counts from IFNβ-treated WT and *Ifnar2* KO HeLa cells 2 hour post Stm infection. Infected cells were treated with gentamicin (Gm) at the indicated concentrations (μg/ml). Data were normalized to the WT+IFNβ Gm 10 group. Mean ± s.d., n = 5. (F) Representative images of Gal3-GFP-expressing HeLa cells 4 hour post mCherry-Stm infection. Scale bar, 10μm. (G) Quantification of the Gal3 positive SCVs from 10 cells. Mean ± s.d., n = 3. Statistical analysis was performed by two-tailed Student’s *t* test (***P < 0.001). See also Figure S5.

The Stm virulence program can lead to the breakage of SCV, exposing the pathogen to the host cytosol (Roy et al., 2004; Xu et al., 2019). To assess whether the pathogen was cytosol-exposed, infected cells were treated with high concentrations of gentamicin, an antibiotic that can penetrate into cells at high concentrations (Myrdal et al., 2005). Stm in IFNβ-treated WT cells were markedly more sensitive to gentamicin than bacteria in IFNβ-treated *Ifnar2* KO cells (Figure 6E), suggesting that IFN-I activation of Stm virulence gene expression promotes SCV rupture and facilitates the pathogen’s access to the cytosol. Consistent with this idea, ∼60% of Stm stained positive for galectin-3, a marker of SCV damage (Thurston et al., 2012), in infected IFNβ-primed WT HeLa cells, whereas <20% of Stm were galectin-3+ in infected *Ifnar2* KO cells (Figures 6F, G). Together, these data suggest a model that explains why IFN-I signaling was a hit in the CRISPR/Cas9 screen: IFN-I signaling-dependent lysosome acidification stimulates intracellular Stm virulence gene expression, which promotes SCV rupture and subsequent Stm*-*induced cytotoxicity.

### IFN-I promotes epithelial Stm pathogenesis *in vivo*

To understand the function of IFN-I signaling in Stm pathogenesis, we used a more physiologic culture system - primary human-derived small intestinal organoids. IFNβ priming of organoids increased cell death associated with Stm infection, whereas treatment of organoids with pyridone-6 had the opposite effect (Figures 7A, B), supporting the idea that IFN-I signaling promotes Stm pathogenicity in IECs.

**Figure 7.**
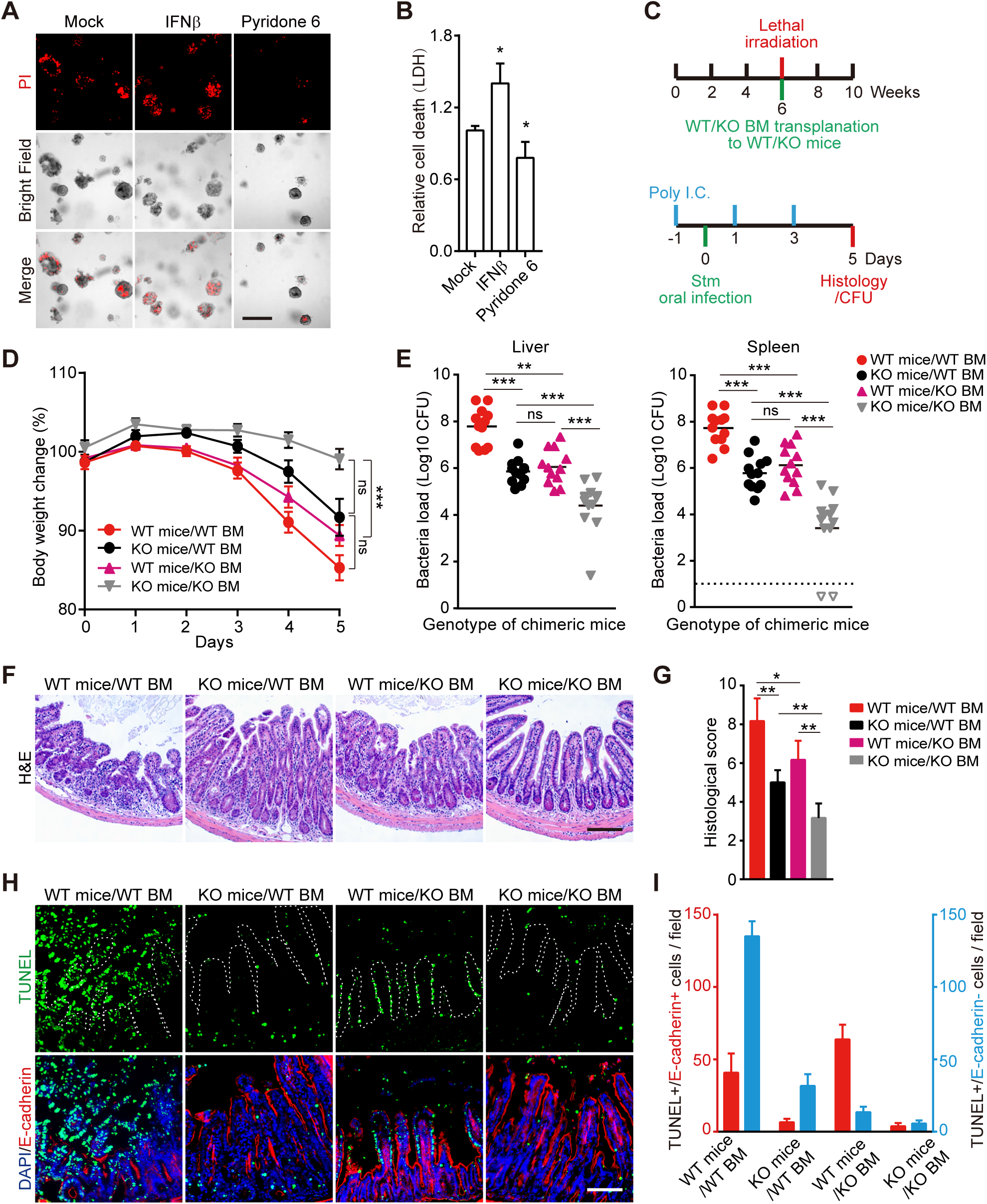
IFN-I signaling in intestinal epithelial cells promotes Stm pathogenesis. (A) Representative images of IFNβ or pyridone-6-primed or unprimed human small bowel enteroids 20 hours post WT Stm infection. Propidium iodide (PI) staining was used to detect cell death. Scale bar, 100Lμm. (B) Enteroid survival 20 hours post WT Stm infection. Mean ± s.d., n = 3. (C) Timeline of generation (top) and oral Stm infection (bottom) of *Ifnar1* chimeric mice. (D) Body weights of Stm*-*infected chimeric mice. Mean ± s.e.m., n = 12 mice. (E) Liver and spleen Stm CFU burdens from chimeric mice 5 days post-Stm infection. Mean ± s.d., n = 12 mice. (F) Representative H&E stained ileal sections from chimeric mice 5 days post-Stm infection. Scale bars, 100μm. (G) Average histological scores of chimeric mice 5 days post-Stm infection from 8 fields. Mean ± s.d., n = 4 mice. (H) Representative images of ileal sections from chimeric mice 5 days post-Stm infection. IECs were identified with E-cadherin (red), dying cells with TUNEL (green), and nuclei with DAPI (blue). The white dashed line marks the epithelial surface. Scale bar, 100μm. (I) Quantification of TUNEL+/E-cadherin+ (red) or TUNEL+/E-cadherin-(blue) cells per field from 8 fields. Mean ± s.d., n = 4 mice. Statistical analysis was performed by two-tailed Student’s *t* test in (B) and (G). Statistical analysis was performed by two-tailed Mann-Whitney U-test in (D) and (E). (*P < 0.05, **P < 0.01 and ***P < 0.001). See also Figure S6.

To further dissect the importance of IFN-I signaling in the context of *in vivo* Stm infection, we used bone marrow transfers to generate chimeric C57BL/6 mice that had *Ifnar1* deleted in only the hematopoietic compartment or in other bodily tissues, including epithelial surfaces (Figure S6A-C). Following intraperitoneal delivery of poly (I:C) to induce IFNβ production (Lauterbach et al., 2010), chimeric mice were oro-gastrically inoculated with Stm to assess the roles of epithelial and hematopoietic compartments in resistance to infection (Figure 7C). Strikingly, mice with KO epithelium and WT bone marrow were relatively resistant to oral Stm infection, with reduced weight loss and distal organ bacterial loads compared to mice that had WT epithelium and bone marrow (Figures 7D, E), suggesting that IEC IFN-I signaling enhances Stm pathogenicity during infection. We also observed a similar phenotype in mice with WT epithelium and KO bone marrow (Figures 7D, E), consistent with previous observations that immune cell IFN-I signaling also promotes Stm pathogenesis (Robinson et al., 2012). Mice that had both KO epithelium and bone marrow were more protected than either chimera (Figures 7D, E), further supporting the idea that Stm takes advantage of IFN-I signaling in both the gut epithelium as well as in bone marrow-derived cells.

Although histological analyses revealed similar levels of tissue damage in both chimeras (Figures 7F, G), finer-scale immunofluorescence studies with TUNEL staining to quantify cell death showed that TUNEL+ (dying) cells tracked with the WT compartment. In chimeric mice with WT epithelium, TUNEL staining primarily co-localized with E-cadherin-positive IECs (Figures 7H, I). In contrast, in Stm*-*infected chimeric mice with WT bone marrow, cell death was primarily localized to E-cadherin-negative cells in the lamina propria (Figures 7H, I). Together, these *in vivo* studies suggest that IFN-I signaling in the epithelial compartment facilitates Stm-induced IEC death and pathogen spread.

## DISCUSSION

Our findings underscore the utility of model intracellular pathogens like Stm as probes for the investigation of fundamental cell processes. The top hits in the genome-scale CRISPR/Cas9 screen that initiated this study were remarkably coherent and revealed that IFN-I signaling sensitizes epithelial cells to Stm cytotoxicity. IFN-I-dependent lysosome acidification in IECs stimulated Stm virulence gene expression, heightened SCV damage and exacerbated cell death, offering a plausible molecular pathway that explains the results of the screen. Importantly, our work uncovered a fundamental new role for IFN-I signaling. We discovered that this canonical antiviral signaling pathway, which has been studied for more than 5 decades (Gonzalez-Navajas et al., 2012), controls the subcellular localization, protein content, pH, and protease activity of lysosomes. Several ISGs, including *Ifitm3, Slc15a3*, and *Cnp*, that were found to localize to lysosomes, were shown to contribute to lysosomal acidification. Thus, IFN-I signaling controls the function of an organelle – the lysosome – in addition to directly or indirectly modulating the expression and activities of hundreds of ISGS and their interaction partners.

The functions of the three ISGs that were identified as participants in IFN-I-mediated lysosomal acidification suggests that more than one mechanism accounts for this phenotype. It seems likely that putative v-ATPase-associated proteins such as *Ifitm3* contribute to acidification processes by directly modulating the proton concentration gradient. However, ISGs with known non-ATPase-related functions such as *Slc15a3*, which is a proton-coupled histidine and di-tripeptide transporter, may not play similar roles. While we cannot exclude v-ATPase-mediated mechanisms for such proteins, we speculate that *Slc15a3* may show transport preferences for non-neutral dipeptides that could influence lysosome luminal pH. Previous studies have linked *Cnp* with not only lysosomal, but mitochondrial compartments (McFerran and Burgoyne, 1997), raising the possibility that ISG function in additional cell compartments could also indirectly contribute to lysosomal acidification. Interestingly, *Cnp*, a membrane-bound protein, has additionally been linked to microtubule function, suggesting that it may also play a role in IFN-I-mediated lysosomal repositioning (Bifulco et al., 2002).

Although a virtually universal antiviral immune signal, the consequences of IFN-I signaling on bacterial pathogens has remained less clear and is in many cases detrimental to the host (Kovarik et al., 2016). Several intracellular bacterial pathogens, including *Listeria monocytogenes, Mycobacterium tuberculosis* and Stm, appear to have decreased virulence in IFNAR1-deficient mice. Studies of the bases for these phenotypes have primarily focused on immune-mediated explanations (Boxx and Cheng, 2016; Kernbauer et al., 2013; O’Connell et al., 2004) (bring back previous citations). For Stm, we propose that IFN-I signaling contributes to the outcome of infection at least in part by inducing remodeling of epithelial cell lysosomes and thereby stimulating the Stm virulence program. Our findings suggest that IFN-I signaling can modify innate defense in the epithelial as well as the immune compartment and is complementary and compatible with previously proposed mechanisms for IFN-I-enhanced Stm infection in bone-marrow derived immune cells. Such mechanisms include elevated macrophage necroptosis (Hos et al., 2017; Robinson et al., 2012) and transcriptional reprogramming (Perkins et al., 2015), altered dendritic cell homeostasis (Stefan et al., 2017), and increased neutrophil-mediated inflammation (Wilson et al., 2019). The role of IFN-I modulation of lysosome function to Stm infection in non-epithelial cells, such as macrophages, requires further study. Furthermore, it remains an open question whether Stm purposely stimulates and exploits IFN-I signaling as part of its pathogenic strategy.

Although IFN-I-induced lysosomal acidification sensitizes cells to an intracellular bacterial pathogen, our finding that several known ISGs with antiviral properties, such as *Ifitm3, Slc15a3* and *Cnp*, participate in this process leads us to speculate that this mechanism may be protective against viral threats. IFN-I-mediated lysosomal remodeling may also play a role in non-infectious pathologies, such as lysosomal cholesterol accumulation (Kuhnl et al., 2018) and other lysosome-related disorders. It remains unclear whether these effects might be driven by the tonic IFN-I signaling that occurs in many tissues (Schoggins et al., 2014), or instead require pathogenic elevations of IFN-I.

Our finding that IFN-I signaling governs the composition and function of the lysosome provides a new cell biologic perspective for understanding cytokine function. It will be of interest to determine whether other immune signals (i.e. including other IFNs and cytokines) can also direct remodeling of lysosomes and other organelles, such as the mitochondria and endoplasmic reticulum, under homeostasis and in diverse pathogenic contexts. Such activities may constitute a broadly applicable lens through which to view and enhance our understanding of the cell biology of innate defense.

## AUTHORCONTRIBUTIONS

H.L.Z. and M.K.W. conceived and all authors designed the study. H.L.Z., A.Z., and X.L. performed all experiments and analyzed data. H.L.Z., B.S., and M.K.W. wrote the manuscript and all authors edited the paper.

## ACKNOWLEDGEMENTS

We thank members of the Waldor lab for helpful discussions on all aspects of this project, Dr. David Breault at the Harvard Digestive Diseases Center (HDDC) Organoid Core for the primary human small intestine organoids; and Michal Pyzik from Dr. Richard Blumberg’s lab for assistance in creation of bone marrow chimeric mice.

Research in the M.K.W. laboratory is supported by HHMI and NIH grant R01 AI-042347. A.Z. was supported by an EMBO long-term fellowship (ALTF 1514-2016) and by a HHMI Fellowship of the Life Sciences Research Foundation.

## DECLARATION OF INTERESTS

The authors declare no competing interests.

**Figure S1. IFN-I promotes intracellular Stm cytotoxicity, Related to Figure 2**

(A) Relative expression of the IFN-I target gene *oas1* in WT and *Ifnar2* KO HT29 cells 16 hours post-IFNβ treatment. Mean ± s.d., n = 3.

(B) Survival of IFNβ-primed or unprimed WT or *Ifnar2* KO HT29 cells 20 hours post WT or mutant Stm infection. Mean ± s.d., n = 3.

(C) Representative bright-field images of WT and *Ifnar2* KO HT29 cells 2 days post Stm infection. Scale bar, 250 μm or 100 μm, respectively.

(D) Quantification of flow cytometry data in Figure 2C. Mean ± s.d., n = 4.

(E) Flow cytometry of IFNβ-primed or unprimed WT and *Ifnar2* KO HeLa cells 20 hours post mCherry-Stm infection and stained with Annexin V-FITC.

(F) Quantification of flow cytometry data from Figure S2E. Mean ± s.d., n = 4.

Statistical analysis was performed by two-tailed Student’s *t* test (*P < 0.05, **P < 0.01 and ***P < 0.001).

**Figure S2. IFN-I does not affect Stm invasion or SCV formation, Related to Figure 2**

(A) Flow cytometry of IFNβ-primed WT and *Ifnar2* KO HT29 cells 4 hours post mCherry-Stm infection. Quantification is shown in Figure 2D.

(B) Representative images of Rab5-RFP-expressing HeLa cells at 4 hours post GFP-Stm infection. Scale bar, 10 μm.

(C) Quantification of Rab5-RFP-positive Stm from 10 fields. Mean ± s.d., n = 3.

(D) Representative images of Rab7-RFP-expressing HeLa cells 4 hours post GFP-Stm infection. Scale bar, 10 μm.

(E) Quantification of Rab7-RFP-positive Stm from 10 fields. Mean ± s.d., n = 3.

**Figure S3. IFN-I signaling regulates lysosomal remodeling in both epithelial cell and THP1 cells, Related to Figure 3**

(A) Quantification of mean fluorescence intensity (MFI) from Figure 3C. Mean ± s.d., n = 3.

(B) Flow cytometry of LysoSensor staining in WT and *Ifnar2* KO HeLa cells ± 16 hours of IFNβ treatment.

(C) Flow cytometry of LysoTracker staining in HT29 cells ± 16 hours of drug treatment.

(D) Quantification of mean fluorescence intensity from Figure 3D. Mean ± s.d., n = 3.

(E) Flow cytometry of Dextran-568 uptake in WT and *Ifnar2* KO HeLa cells ± 16 hours of IFNβ treatment.

(F) Flow cytometry of LysoTracker staining in monocytic macrophage-like THP1 cells ± 16 hours of IFNβ treatment.

Statistical analysis was performed by two-tailed Student’s *t* test (**P < 0.01 and ***P < 0.001).

**Figure S4. Purity of isolated lysosomes and IFITM3 gene KO in HeLa cells, Related to Figure 5**

Immunoblotting for protein markers of indicated subcellular compartments in whole-cell lysates (T) and purified lysosomes (IP).

**Figure S5. Intracellular Stm virulence gene expression, Related to Figure 6**

(A-D) Relative induction of individual SPI-2 (A), PhoP-induced (B), SPI-3 (C) or SPI-1 (D) genes in intracellular Stm from WT and *Ifnar2* KO HeLa cells ± drug treatment. Data are normalized to transcript levels from LB-cultured Stm (red). Mean ± s.d., n = 3. Statistical analysis was performed by two-tailed Student’s *t* test (***P < 0.001).

**Figure S6. Generation of chimeric mice by bone marrow transfer, Related to Figure 7**

(A-C) Flow cytometry of peripheral blood CD45.1 and CD45.2+ cells in mock and chimeric mice 4 weeks after bone marrow transplantation. WT mock mice carry CD45.2 allele but not CD45.1 (A), which is abolished by irradiation (B, C). 4 weeks later after CD45.1 BM transfer, the chimeric mice carry CD45.1 allele but not CD45.2 (B, C).

## STAR METHODS

### KEY RESOURCE TABLE

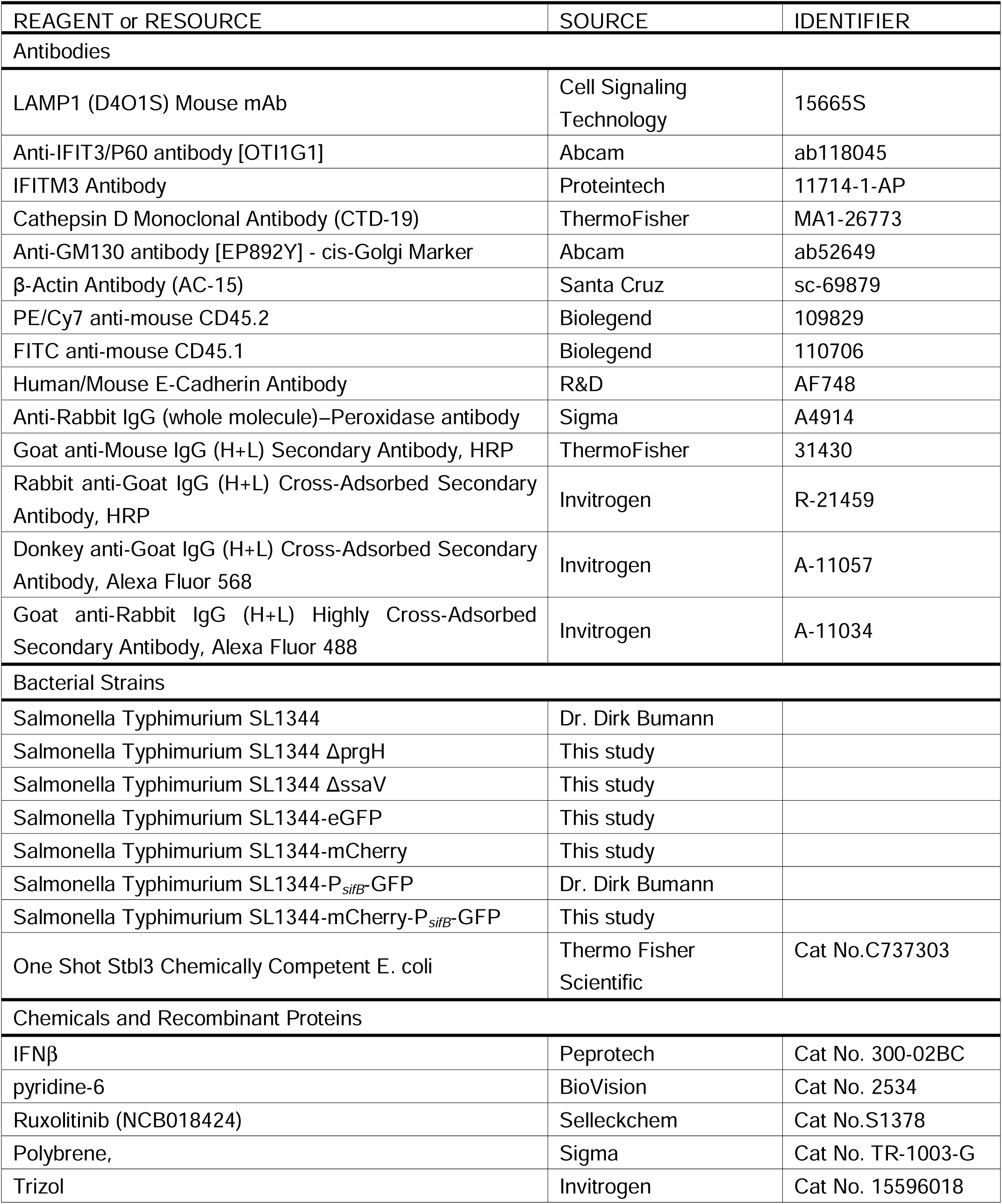
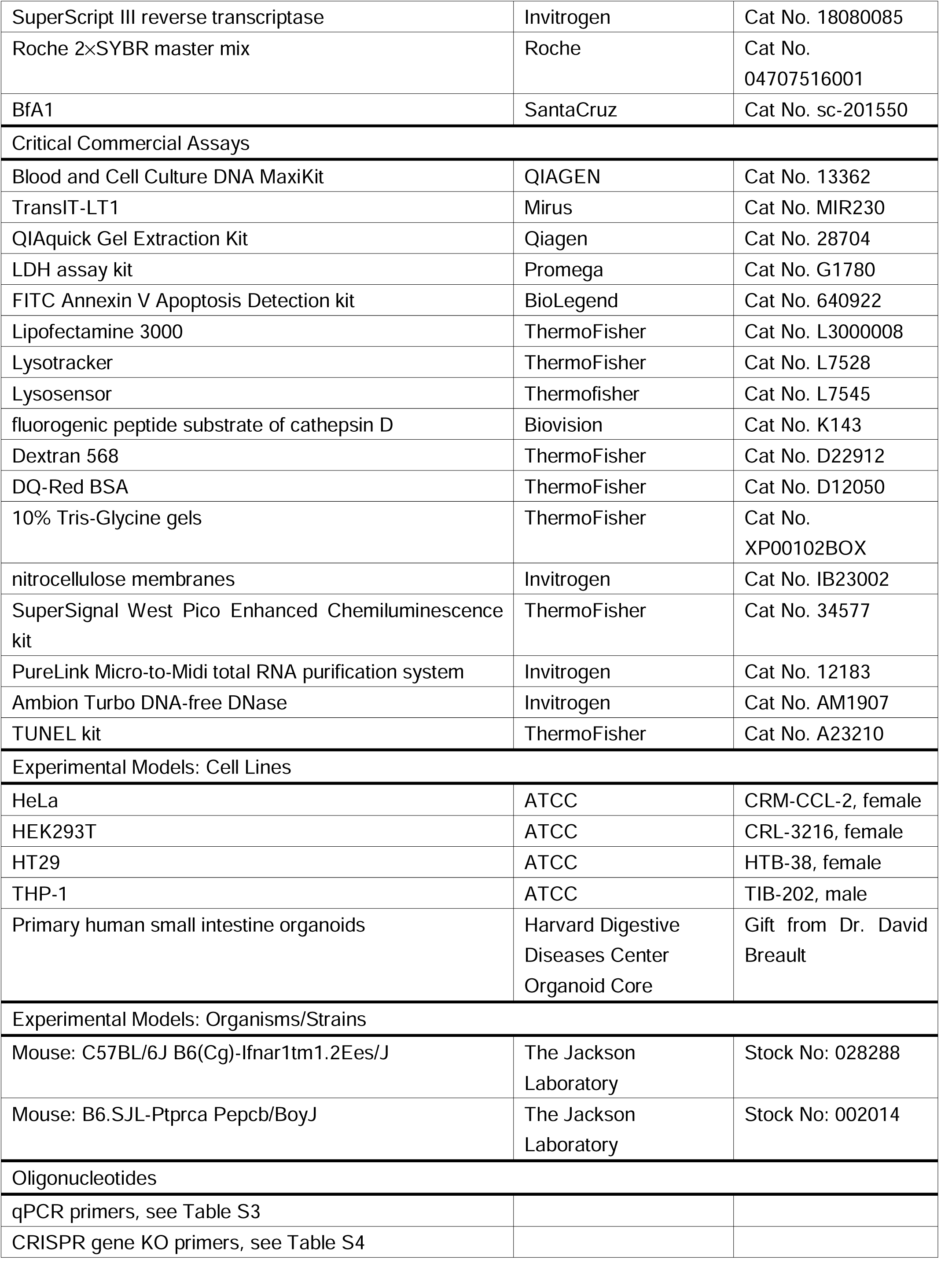
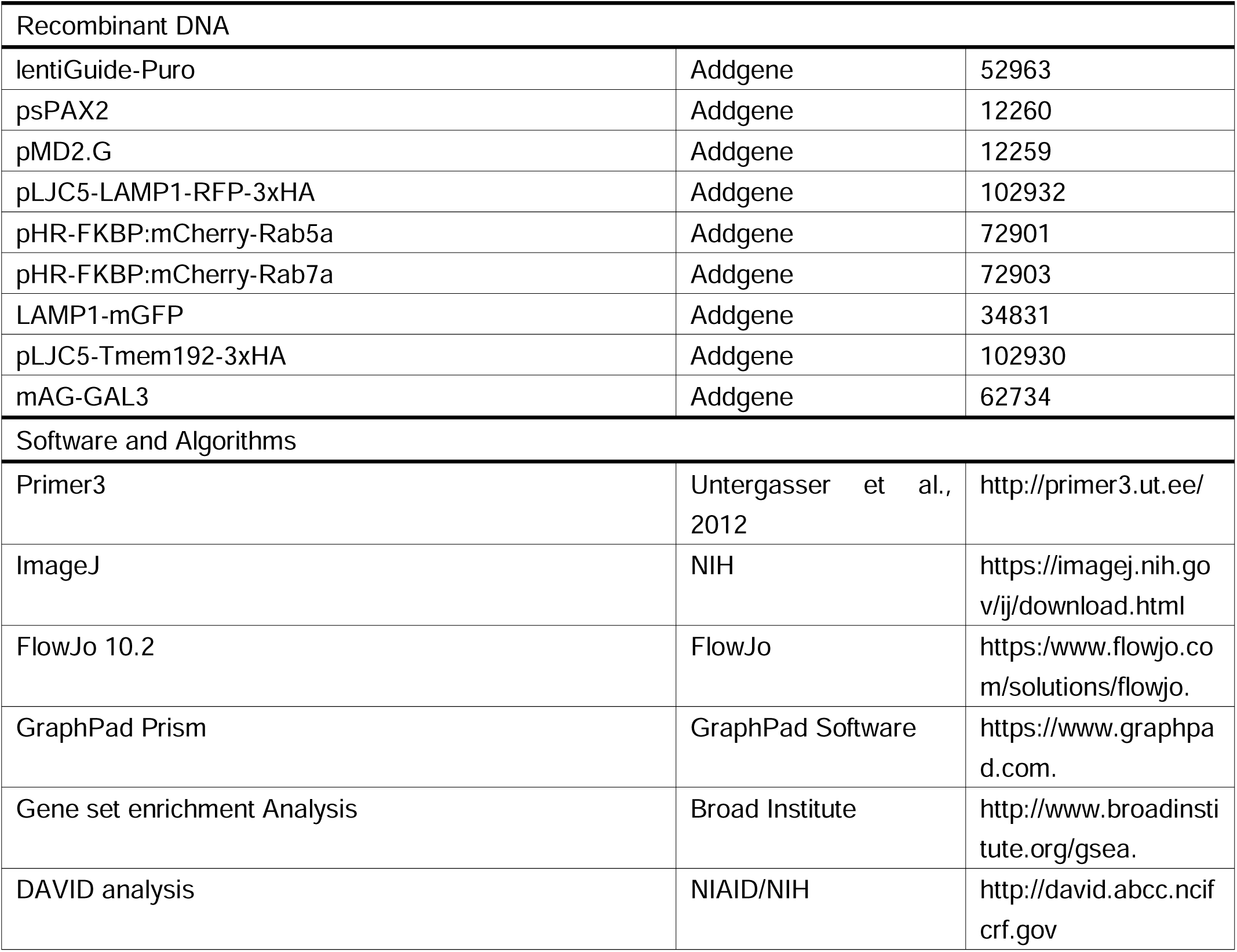

## CONTACT FOR REAGENT AND RESOURCE SHARING

Further information and requests for resources and reagents should be directed to and will be fulfilled by the Lead Contact, Matthew K Waldor

## EXPERIMENTAL MODEL AND SUBJECT DETAILS

### Bacterial strains, plasmids, and antibodies

Strains, plasmids, oligonucleotides and antibodies used in this study are listed in key resources table and table S3, 4. *Escherichia coli* K-12 DH5α λ pir was used for cloning procedures and plasmid propagation. *S. typhimurium* strain SL1344 and its ΔSPI-1 and ΔSPI-2 derivatives were cultured in Luria-Bertani (LB) medium or on LB agar plates at 37°C supplemented with streptomycin (100μg/ml). The SPI-1 (*prgH*) and SPI-2 (*ssaV*) genes were deleted from wild-type (WT) SL1344 with the lambda red recombination system (Datsenko and Wanner, 2000). This approach was also used to introduce the GFP and mCherry-coding sequence with a constitutive promoter (P_*rpsM*_) into the *putP-putA* locus (Hautefort et al., 2003).

### Cell lines

HeLa (ATCC, Cat No. CRM-CCL-2, female) and HEK293T (ATCC, Cat No. CRL-3216, female) cells were cultured in Dulbecco’s modified Eagle’s medium (DMEM; ThermoFisher, Cat No. 11965) supplemented with 10% fetal bovine serum (FBS; Gibco, Cat No. 16140-071). HT29 (ATCC, Cat No. HTB-38, female) cells were cultured in McCoy’s 5A modified medium (Thermo Fisher, Cat No. 30-2007) supplemented with 10% FBS. THP-1 (ATCC, Cat No. TIB-202, male) cells were cultured in RPMI-1640 medium (Lonza, Cat No. 12-167F) with 10% non-heat inactivated FBS (GeminiBio, Cat NO. 100-500) and supplemented with HEPES (Lonza, Cat No. 17-737E), 2-Mercaptoethanol (Invitrogen, Cat No. 21985023) and L-Glutamine (Lonza, Cat No. 17-605E). All cells were cultured at 37°C in a 5% CO_2_ incubator.

### Infection of organoids derived from human small intestine

Primary human small intestine organoids (enteroids) were kindly provided by Dr. David Breault at the Harvard Digestive Diseases Center (HDDC) Organoid Core. Enteroids were cultured in the following medium: advanced DMEM/F12 (Gibco, Cat No.12634-028) supplemented with L-WRN conditioned medium (ATCC CRL-3276; HDDC Organoids Core), HEPES (10mM, pH 7.4), GlutaMax (Gibco, Cat No.35050-061), B_27_ (Gibco, Cat No.12587010), N2 (Gibco, Cat No.17502-048), 1mM N-acetyl-L-cysteine (Sigma, Cat No.A8199), 10mM nicotinamide (Sigma, Cat No.N0636), 5µM A83-01 (Sigma, Cat No.SML0788), 10µM SB202190 (Sigma, Cat No.S7067), 50ng/ml murine EGF (Peprotech, Cat No.315-09), 10nM gastrin (Sigma, Cat No.G9145), and 10µM Y-27632 (Sigma, Cat No.Y0503). For Stm infection, approximately 100 enteroids were seeded in 50µl Matrigel (Corning, Cat No.356231) in each well of a 24-well plate. Three to four days after seeding, enteroids were either mock treated, or primed with either 10ng/ml IFNβ or 0.5 µM pyridine-6 for 16 hours. Enteroids were then released from Matrigel by incubation in 500µl Cell Recovery Solution (Corning, Cat No.354253) for 30 mins on ice. Resuspended enteroids were pipetted up and down 50 times with a P1000 pipette and then transferred to a new 24-well plate. Each well was infected with approximately 3×10^7^ Stm. The plate was centrifuged for 5 mins at 300× g before it was placed in a 37°Cincubator for 30 mins. After infection, enteroids were transferred to microcentrifuge tubes, spundown, mixed with 50µl Matrigel per tube/sample, and seeded into a new 24-well plate. After Matrigel solidification at 37°C, 500µl full enteroid media containing 50μg/ml gentamicin was added to each well at 10ng/ml IFNβ or 0.5µM pyridine-6 was added to the corresponding primed samples. At 20 hpi, propidium iodide (Invitrogen, Cat No.V13241) was used to stain the enteroids before imaging. To quantitatively measure cell death, the media from each well/sample was also assayed for LDH activity as described above. LDH release values of mock treated samples were set at 1 for normalization.

### Bone marrow chimera mice

C57BL/6 and *Ifnar1*^*-/-*^ mice were purchased from The Jackson Laboratory (Bar Harbor, ME, USA) and were maintained on a 12-hour light/dark cycle and a standard chow diet at the Harvard Institute of Medicine specific pathogen-free (SPF) animal facility (Boston, MA, USA). Animal experiments were performed according to guidelines from the Center for Animal Resources and Comparative Medicine at Harvard Medical School. All protocols and experimental plans were approved by the Brigham and Women’s Hospital Institutional Animal Care and Use Committee (Protocol #2016N000416). Littermate control male and female mice were randomly assigned to each group and experiments were performed blinded with respect to treatment. For bone marrow chimeras, recipient mice were irradiated two times with 600 rad 1□day before injection of bone marrow from WT or *Ifnar1*^-/-^ mice. Bone marrow was extracted from femurs of donor mice by flushing with PBS and then washed once in PBS; 1× 10^6^ cells were injected into the tail vein of recipient mice. Mice were monitored for 4 weeks, at which point engraftment was evaluated by flow cytometry.

### Infection of chimeric mice

20μg poly (I:C) (Sigma, Cat No. P1530) was given intraperitoneally to chimeric mice one day before Stm infection and every other day for a total of 3 doses to stimulate IFN production. Food was withdrawn for 4 hours before infection. Stm inocula were prepared as described above. Mice were infected orogastrically with 5□×□10^8^ Stm in 100μl PBS. Food was returned to the cages 2hpi. Infected mice were sacrificed 5 days after infection. Tissue samples of the small intestine, spleen and liver were collected for histological analysis and enumeration of colony-forming units (CFU). CFU were quantified by serial-dilution plating of homogenized tissue samples on LB plates containing 100μg/ml streptomycin.

## METHOD DETAILS

### Pharmacologic inhibitors and IFNβ priming

JAK inhibitors pyridine-6 (BioVision, Cat No. 2534) and ruxolitinib (NCB018424) (Selleckchem, Cat No.S1378) were used at 0.5µM. IFNβ (Peprotech, Cat No. 300-02BC) was used at 10ng/ml for cell priming. Drug-treated cells were primed for 16 hours (unless otherwise indicated) before Stm infection.

### Stm infections

All tissue culture infections were done according to the following procedure unless otherwise indicated. WT and mutant Stm were grown for ∼16 hours at 37°C with shaking and then sub-cultured (1:33) in LB without antibiotics for 3 hours until the cultures reached an OD_600_ of 0.8.To prepare the inoculum, cultures were first pelleted at 5,000× g for 5 min. The pellets were resuspended in DMEM without FBS, and an appropriate volume of bacterial solution was added to cells to reach a multiplicity of infection (MOI) of 100 bacteria per eukaryotic cell. The cells were then incubated with bacteria for 30 min at 37°C with 5% CO_2_. Extracellular bacteria were removed by extensive washing with phosphate-buffered saline (PBS; Gibco, Cat No. 14190250) and addition of 50μg/ml gentamicin to the medium. At 2 hours post infection (hpi), the gentamicin concentration was decreased to 5µg/ml.

### CRISPR/Cas9 Stm infection screen

HT29-Cas9 CRISPR libraries were constructed as described previously (Blondel et al., 2016) using the Avana sgRNA library, which contains four different sgRNAs targeting each human protein-coding gene (Doench et al., 2016). For each library, two sets of four T225 flasks (Corning, Cat No. 14-826-80) were seeded with 15 × 10^6^ cells per flask and then incubated for 48 hours. At the time of the screen, there were ∼150× 10^6^ cells per experimental condition, corresponding to ∼2,000× coverage per sgRNA. Cells were at ∼70% confluence at the time of infection. The infection was done as described above with minor modifications. Briefly, HT29 libraries were infected with WT Stm at an MOI of 300 for 30 min. After infection, the libraries were expanded in McCoy’s 5A + FBS containing 5µg/ml gentamicin, to both permit intracellular bacterial cytotoxicity and minimizethe intracellular gentamicin concentration to allow Stm invasion during the next round of infection. Flasks were checked daily to monitor recovery of survivor cells; when 70% confluency was achieved, cells were trypsinized, pooled, and reseeded for the next round of infection. In total, four rounds of infection were conducted. Surviving cells from the last round of infection were used for preparation of genomic DNA.

### Genomic DNA preparation, sequencing, and analyses of screen results

Genomic DNA was obtained from 75× 10^6^ cells after positive selection, as well as from input cells, using the Blood and Cell Culture DNA MaxiKit (QIAGEN, Cat No. 13362). sgRNA sequences was amplified by PCR as described (Doench et al., 2016). The read counts were first normalized to reads per million within each condition by the following formula: reads per sgRNA / total reads per condition × 10^6^. Reads per million were then log_2_-transformedby first adding 1 to all values, in order calculate the log of sgRNAs with zero reads. The log_2_ fold-change of each sgRNA was then determined relative to the input sample for each biological replicate. MAGeCK analysis for genome-scale CRISPR-Cas9 knockout screens was used to evaluate the rank and statistical significance of perturbations from the ranked list (Li et al., 2014) and enriched pathways were determined using ClueGo (Bindea et al., 2009).

### Lentivirus preparation and transductions

The Galectin 3, Rab5, Rab7, LAMP1, and LC3B lentiviral expression plasmids used in the study are listed in Table S4. Lentiviral packaging plasmids psPAX2 and pVSV-G, and the corresponding cargo plasmid were transfected into 293T cells with the TransIT-LT1 transfection reagent (Mirus, Cat No. MIR230). 48 hours following transfection, 293T culture supernatants were harvested, passed through a 0.45µm pore filter, and added to target cells that were grown to 70-80% confluency in 6-well plates. Polybrene (Sigma, Cat No. TR-1003-G) (8µg/ml) was added and the 6-well plates were spun at 1000×g for 2 hours at 30°C, after which cells were returned to 37°C. The infections were repeated the next day with supernatants from 72 hour-transfected 293T cultures.

### Construction of cell lines with targeted gene disruptions

The sgRNA sequences used for construction of targeted HT29-Cas9 and HeLa-Cas9 mutant cell lines are listed in Table S4. All sgRNA oligonucleotides were obtained from Integrated DNA Technologies (IDT) and cloned into the pLentiGuide-Puro plasmid. Briefly, 5µg of plasmid pLentiGuide-Puro was digested with BmsBI (Fermentas, Cat No. ER0451) and purified using the QIAquick Gel Extraction Kit (Qiagen, Cat No. 28704). Each pair of oligos was annealed and phosphorylated with T4 PNK (NEB, Cat No. M0201S) in the presence of 10× T4 DNA ligase buffer in a thermocycler with the following parameters: i) incubation for 30 minutes at 37°C, ii) incubation at 95°C for 5 min with a ramp down to 25°C at 5°C per minute. Oligos were then diluted 1:200, and 1µl of the diluted oligo mixture was ligated with 50ng of BsmBI digested plasmid. Ligations were transformed into the STBL3 *Escherichia coli* strain (Thermo Fisher, Cat No. C7373-03) and positive clones were identified by Sanger sequencing (Genewiz). Lentiviral transduction of sgRNAs cloned into pLentiGuide-Puro into HT29-Cas9 and HeLa-Cas9 cells was performed as described above. Targeted gene KO cell lines were selected by puromycin (1µg/ml) for 10 days. HT29 KO cells were isolated as single clones while HeLa cells were CRISPR KO pools after drug selection.

### Cell survival assays

For cell survival assays, 5 × 10^4^ HT29 cells were seeded into 96-well plates and primed with or without drugs in McCoy’s 5Amedium supplemented with 10% FBS.HT29 cells were infected with Stms trains at an MOI of 100 as described above. Cell survival analysis was performed using an LDH assay (Promega, Cat No. G1780) according to the manufacturer’s protocol at 4 and 20 hpi.

### Stm invasion assays

mCherry-or GFP-tagged Stm were used in all flow cytometry and immunofluorescence experiments. Stm infections were performed as described above with varying MOIs. At 4hpi, suspended and attached cells were collected, resuspended in PBS, and immediately analyzed with a LSR II (BD Bioscience) or SH800 (Sony) flow cytometer. Data were processed with FlowJo software (v10.6.1).

### Annexin V staining and FACS analysis

Cell death was detected with the FITC Annexin V Apoptosis Detection kit (BioLegend, Cat No. 640922). Infections were performed as described above with mCherry-Stm at an MOI of 100. 20 hpi suspended and attached cells were collected, resuspended in 100μl of Annexin V binding buffer at 1×10^7^ cells/ml and mixed with 5μl of FITC-conjugated Annexin V. After incubation at room temperature (RT) for 15□min in the dark, 400μl of Annexin V binding buffer was added and stained cells were immediately analyzed by flow cytometry as described above.

### Immunofluorescence microscopy of tissue cultured cells

HeLa cells were seeded in 12-well plates on 18 mm glass coverslips or 6-well chambers (Mat-TEK, Cat No.P06G-1.5-10-F). Cells were transiently transfected with LAMP1-GFP expressing plasmid mixed with Lipofectamine 3000 (ThermoFisher, Cat No. L3000008) according to the manufacturer’s instructions. 24 hours post-transfection cells were primed with or without 10ng/ml IFNβ for 16 hours.The cells were then stained with 75nM Lysotracker (ThermoFisher, Cat No. L7528) for 15 min and then fixed with 2% PFA for 20 min at RT. The samples were washed with PBS three times, and stained with fluorescent phalloidin (1:1000) and 4,6-diamidino-2-phenylindole (DAPI, 1□μg□/ml) to label actin filaments and nuclei, respectively. For experiments with LAMP1-RFP, Rab5-RFP, Rab7-RFP, Gal3-GFP, and eGFP-LC3B cell lines, cells were seeded in 6-well chambers and primed with 10ng/ml IFNβ for 16 hours before infection with fluorescently-labeled Stm at an MOI 50. Live cells were analyzed at 2 hpi by confocal microscopy to detect localization of Gal3 and Stm.

### Quantification of lysosome distribution

Lysosome distribution was analyzed as described (Li et al., 2016); the area occupied by nuclei was excluded from analyses. Average LAMP1 intensities were measured for the area within 5μm of the nucleus (*I*_perinuclear_), and the area >10μm from the nucleus (*I*_peripheral_). The average intensities were calculated and normalized to cell areas. The perinuclear index was defined as *I*_perinuclear_ /*I*_peripheral_. Quantifications were carried out on10 cells per group with ImageJ.

### Measurement of lysosome acidity

Cells with no treatment or with either 10ng/ml IFNβ or 5nM BfA1 (SantaCruz, Cat No. sc-201550) treatment for 16 hours were stained with 75nM Lysotracker or Lysosensor (Thermofisher, Cat No. L7545) for 15 min and washed with PBS. The fluorescence intensity of the stained cells was determined by flow cytometry.

### Cathepsin D activity assay

HeLa cells were seeded in 96-well plates with or without 10ng/ml IFNβ priming for 16 hours. A fluorogenic peptide substrate of cathepsin D, Mca-P-L-G-L-Dpa-A-R-NH2 (Biovision, Cat No. K143), was added to the cells to a final concentration of 200μM for 2 hours. The fluorescence intensity of each well was measured with a fluorescence plate reader. Each sample was assayed in triplicate and normalized to a standard curve.

### Endocytosis and lysosome function assays

HeLa cells were seeded in 24-well plates with or without 10ng/ml IFNβ priming. Cells were treated with either 50μg/ml Dextran 568 (ThermoFisher, Cat No. D22912) or 25μg/ml DQ-Red BSA (ThermoFisher, Cat No. D12050) for 2 hours in growth medium. Then, cells were washed with PBS and trypsinized for fluorescence quantification by flow cytometry.

### Lysosome immunopurification (LysoIP)

LysoIP was performed largely as described (Abu-Remaileh et al., 2017). Briefly, pLJC5-3×HA-TMEM192 was used to introduce a lysosomal tag protein in WT and *Ifnar2* KO HeLa cells. 15 million cells were used for each replicate. Cells were rinsed twice with pre-chilled PBS and then scraped in 1ml of PBS containing protease and phosphatase inhibitors and pelleted at 100×g for 2 min at 4°C. Cells were resuspended in 950µl of the same buffer, and 25µl (equivalent to 2.5% of the total number cells) was reserved for further processing to generate the whole-cell sample. The remaining cells were gently homogenized with 25 strokes of a 2ml Dounce-type homogenizer. The homogenate was then centrifuged at 100×g for 2 min at 4°C to pellet the cell debris and intact cells, while cellular organelles including lysosomes remained in the supernatant. The supernatant was incubated with 150µl of anti-HA magnetic beads preequilibrated with PBS on a rotator shaker for 3 min. Immunoprecipitates were then gently washed three times with PBS on a DynaMag Spin Magnet. Beads with bound lysosomes were resuspended in 100µl pre-chilled 1% Triton-X lysis buffer to extract proteins. After 10 min incubation on ice, the beads were removed with the magnet. 5µl of each sample were subjected to 12.5%-acrylamide SDS-PAGE and immunodetected using antibody listed in Table S6, while the remainder was submitted to the Thermo Fisher Center for Multiplexed Proteomics of Harvard Medical School (Boston, MA, USA) for Isobaric Tandem Mass Tag (TMT)-based quantitative proteomics.

### Immunoblot analyses

Mammalian cell lysates were prepared in radioimmuno-precipitation assay (RIPA) buffer supplemented with 1 tablet of EDTA-free protease inhibitor (Roche, Cat No. C762Q77) per 25ml buffer. Lysates were kept at 4°C for 30 min and then clarified by centrifugation in a microcentrifuge at 13,000 rpm at 4°C for 10 min. Proteins were denatured by the addition of SDS sample buffer and boiling for 5 min. Proteins were separated by electrophoresis in 10% Tris-Glycine gels (ThermoFisher, Cat No. XP00102BOX), and then transferred onto nitrocellulose membranes (Invitrogen, Cat No. IB23002). The antibodies and dilutions used are listed in Table S6. Blots were developed with the SuperSignal West Pico Enhanced Chemiluminescence kit (ThermoFisher, Cat No. 34577), and imaged with a Chemidoc (Bio-Rad).

### qRT-PCR quantification of Stm virulence gene expression

Hela cells were seeded at 1.5×10^6^ cells per 6-well plates. After drug-treatment for 16 hours, cells were infected with Stm at an MOI of 50 as described above. Cells were washed with PBS and lysed in Trizol (Invitrogen, Cat No. 15596018) at 1and 4 hpi. RNA was purified with the PureLink Micro-to-Midi total RNA purification system (Invitrogen, Cat No. 12183) according to the manufacturer’s instructions. RNA samples were treated for residual DNA contamination using Ambion Turbo DNA-free DNase (Invitrogen, Cat No. AM1907). Purified RNA was quantified on a Nanodrop 1000 (Thermo Scientific). RNA was reverse transcribed for quantitative RT-PCR (qRT-PCR) experiments by adding 10µg of total RNA to a mixture containing random hexamers (Life Technologies), 0.01M dithiothreitol, 25 mM dNTPs (Thermo Scientific, Cat No. R0191), reaction buffer and 200 units of SuperScript III reverse transcriptase (Invitrogen, Cat No. 18080085). cDNA was diluted 1:50 in dH_2_O and mixed with an equal volume of target-specific primers and Roche 2×SYBR master mix (Roche, Cat No.04707516001). Plates were centrifuged at 1000 rpm for 1 min and stored at 4°C in the dark until ready for use. Primer pairs were designed to minimize secondary structures, a length of ∼20-nucelotides and a melting temperature of 60°C using the primer design software Primer 3. Primer sequences are listed in Table S3. For data normalization, quadruplicate C_t_ values for each sample were averaged and normalized to C_t_ values of the control gene *rpoB*. The relative gene expression level of Stm in infection conditions was normalized to LB-cultured Stm.

### Flow cytometric analysis of Stm virulence gene expression

HeLa cells were infected with mCherry-and *sifB*-GFP-expressing-Stm as described above. Cell lysis was performed 4 hpi by washing three times with PBS and subsequent incubation for 10 min with PBS containing 0.1% Triton X-100. Cell lysates were then analyzed by flow cytometry. Stm were first identified by gating on the mCherry signal and *sifB* expression was quantified by gating on the mCherry+/GFP+ population. LB cultured Stm served as negative control.

### Gentamicin protection assay

Gentamicin protection assays were carried out as described (Knodler et al., 2014). Briefly, HeLa cells in 96-well plates were infected in triplicate with Stm at an MOI of 50. Cells were washed three times with PBS and incubated in medium containing 100µg/ml gentamicin for 30 min to eliminate extracellular bacteria. Then, media with either 10, 100 or 400µg/ml gentamicin was applied to the cells. Cells were lysed 2hpi by washing three times with PBS and subsequent incubation for 10 min with PBS containing 0.1 % Triton X-100. Colony forming units (CFUs) were enumerated by plating serial dilutions of the lysates onto LB plates with 100μg/ml streptomycin. Data was normalized to the CFU of WT HeLa cells at gentamicin 10.

### Histology and tissue immunofluorescence

Formalin-fixed, paraffin-embedded distal small intestinal samples sections of 4μm thickness were mounted on glass slides and stained with hematoxylin and eosin. Histology score was evaluated as described (Erben et al., 2014). For immunofluorescence analysis, distal small intestinal samples were collected and flushed with PBS and fixed in 4% paraformaldehyde (PFA) overnight at 4°C followed by washing with PBS. Tissues were embedded in Optimal Cutting Temperature Compound (Tissue-Tek) and stored at -80°C before sectioning on a CM1860 UV cryostat (Leica). 6μm-thick slides were stained with TUNEL (ThermoFisher, Cat No. A23210) according to the manufacturer’s instructions and then incubated with anti-E-cadherin antibodies at 4°C overnight at a 1:200 in PBS. The next day, AF568-conjugated secondary antibody, diluted at 1:500, was applied to the slides for 1 hour. Nuclei were stained with DAPI at RT for 5 min in the dark. Samples were imaged with an Eclipse Ti confocal microscope with a 20×□objective (Nikon).

## QUANTIFICATION AND STATISTICAL ANALYSIS

Statistical analyses were carried out using the two-tailed Student’s *t* test or one-way analysis of variance (ANOVA) with Dunnet’s post-correction on GraphPad Prism5.

## DATA AND CODE AVAILABILITY

Original data for results of CRISPR screening is in Table S1, and original data for mass spectrometry of lysosome proteomic is in Table S2.

### Supplementary items

Table S1: CRISPR/Cas9 screening results, related to figure 1

Table S2: Mass spectrometry of lysosome proteomic, related to figure 3

Table S3: qPCR primers

Table S4: CRISPR KO primers

