## Supplementary figures and images for "Functional remodeling of lysosomes by type I interferon modifies host defense"

### Supplemental figures

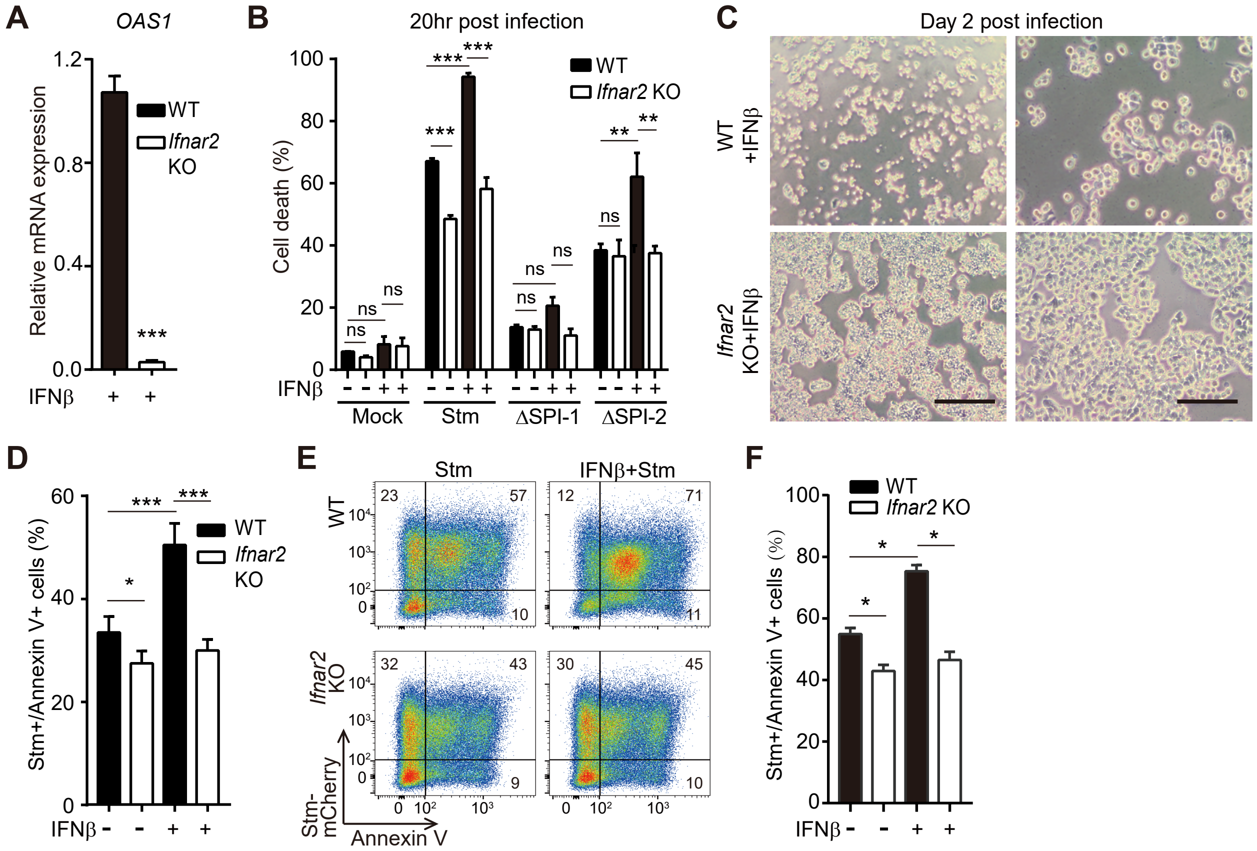

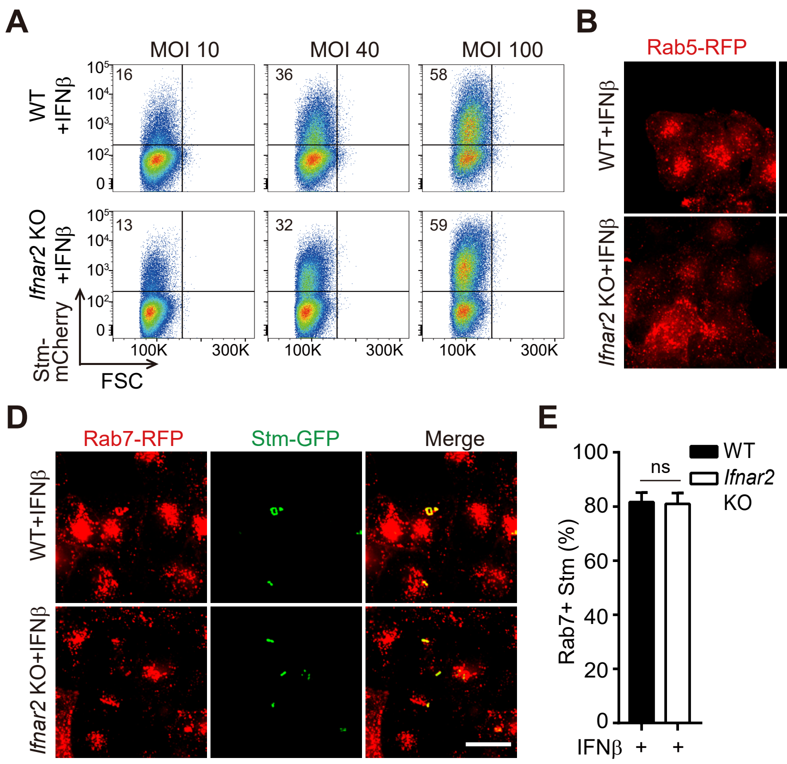

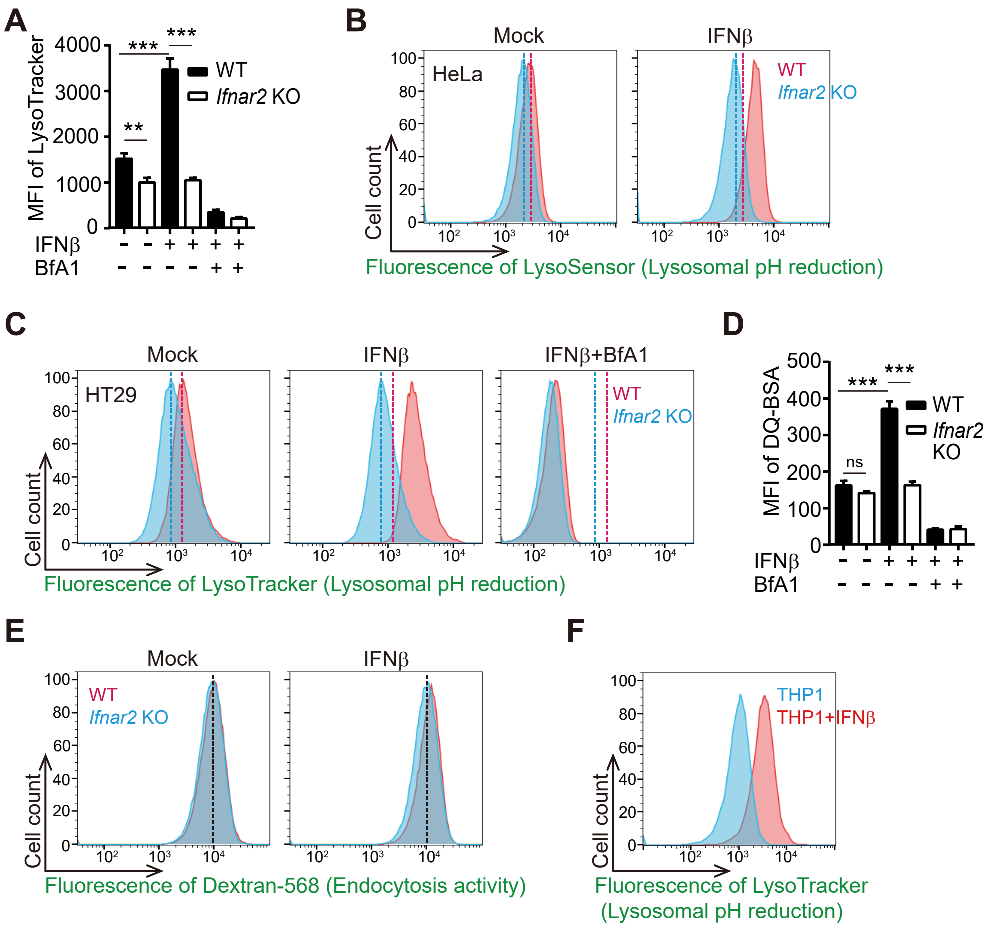

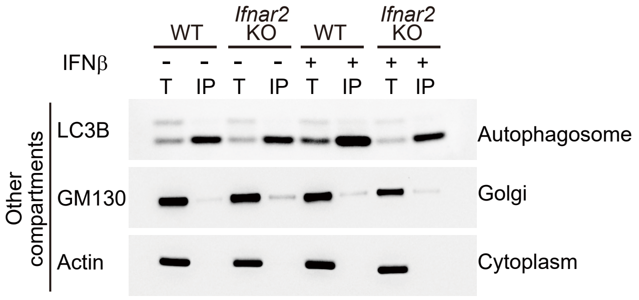

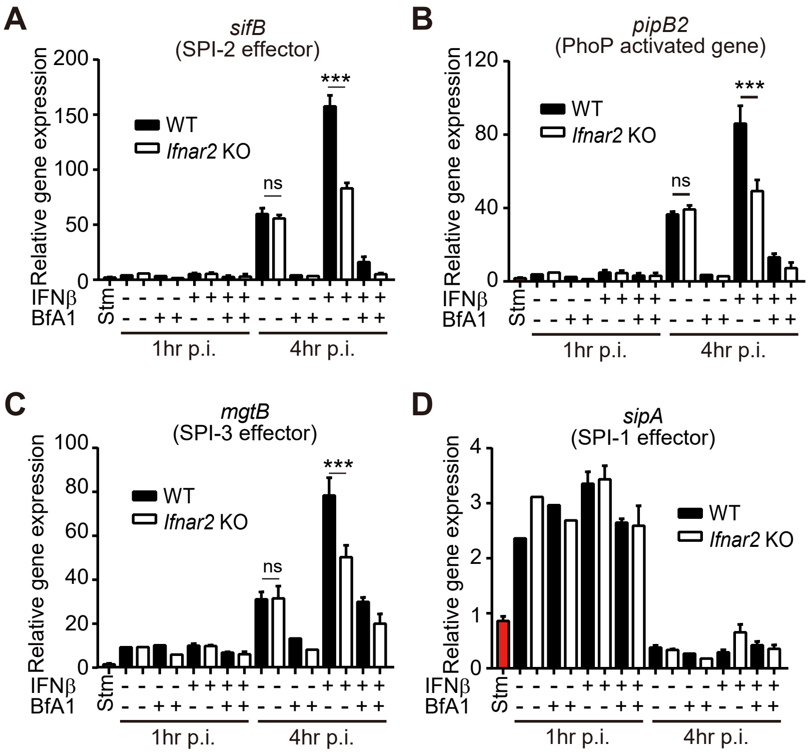

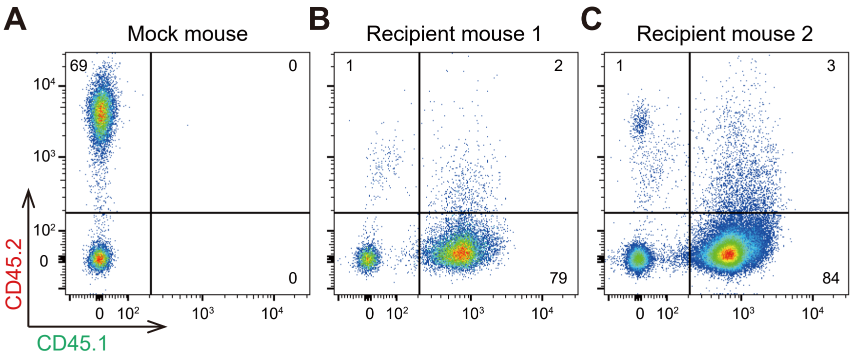
